# Transcriptomic Profiling and Regulatory Network Analysis of Ten Metabolic Transporters Across Five Diabetic Complications: A Multi-Dataset, Twelve-Phase GEO Bioinformatics Study

**DOI:** 10.64898/2026.05.23.727195

**Authors:** Babatunde B. Adegboyega, Philip E. Courage, Olamide O. Awolaja, Elaho Osarietin, Benson Okorie

**Author notes:** 267.

## Abstract

**Objective:** Diabetic complications collectively represent one of the most urgent unresolved problems in medicine, yet the field continues to study them in near-complete isolation from one another. No unified framework has systematically characterised the shared and divergent molecular signatures of ten clinically critical metabolic transporters across all five major complications, cardiomyopathy (DCM), nephropathy (DN), retinopathy (DR), peripheral neuropathy (DPN), and atherosclerosis and vasculopathy (DAD), through an integrated, multi-method computational pipeline. This study was designed to address that gap directly.

**Methods:** Eleven GEO microarray datasets comprising 118 diabetic and 76 control samples were analysed through twelve sequential phases: differential expression analysis, pan-complication overlap, weighted gene co-expression network analysis (WGCNA), GO/KEGG functional enrichment with gene set enrichment analysis (GSEA), STRING protein-protein interaction (PPI) network construction, competing endogenous RNA (ceRNA) network mapping, transcription factor activity inference using a VIPER-style algorithm, immune cell infiltration estimation by single-sample GSEA, diagnostic biomarker modelling using LASSO logistic regression and Random Forest classification, CMap-style drug repurposing by connectivity scoring, and two-sample Mendelian randomisation (MR) employing four independent estimators (inverse-variance weighted [IVW], MR-Egger, weighted median, and weighted mode).

**Results:** CD36 was the only transporter to achieve significant dysregulation across three independently sourced tissue types (DN, DR, DPN; logFC range 0.88 to 2.18), whilst TLR4 exhibited the highest fold-change in the study (logFC = 3.88, DPN) and the greatest WGCNA module membership (kME = 0.976, DPN). SERCA2 was significantly downregulated in three complications (DCM, DN, and DR) at formal significance thresholds and trended negatively in the remaining two (DPN and DAD), constituting the most consistently suppressed transporter in the study. Its universal downregulation was explicable through four convergent mechanisms spanning transcriptional, oxidative, ceRNA-mediated, and transcription factor-level regulation, and was confirmed as causally relevant to diabetic cardiomyopathy by eQTL Mendelian randomisation (beta = -0.085, p = 0.005). miR-21-5p was identified as the dominant ceRNA regulatory bridge (betweenness centrality = 0.428; 6.7-fold above the second-ranked miRNA), with MALAT1 as the sole lncRNA hub active in all five complications. PPARgamma and TP53 repression emerged as the leading transcription factor-level explanations for the simultaneous metabolic and inflammatory dysregulation characteristic of the diabetic transcriptome. Immune deconvolution revealed DCM as immunologically quiescent, DN as comprehensively infiltrated (ten enriched cell types), and DPN as mast-cell-dominated, identifying a cellular mechanism for TLR4-driven neuroinflammation that has not previously been systematically characterised. GLUT4 achieved perfect diagnostic discrimination for DPN (AUC = 1.000, p < 0.001; LASSO coefficient = -2.143), whilst SGLT2 was the leading DAD diagnostic marker (AUC = 1.000, p = 0.002). Epalrestat was the sole pan-complication drug repurposing candidate (significant connectivity reversal in four of five complications). Mendelian randomisation confirmed causal effects of T2DM genetic liability on all five complications (all p < 0.0001, all four estimators concordant), and eQTL-MR identified TLR4 (beta = +0.073, p = 0.006) and CD36 (beta = +0.070, p = 0.008) as causal risk factors for DN, SERCA2 reduced expression as a causal driver of DCM (beta = -0.085, p = 0.005), and SGLT2 expression as a causal protector against DN (beta = -0.070, p = 0.013).

**Conclusions:** This twelve-phase investigation identifies a pan-complication CD36/TLR4 inflammatory dyad and a SERCA2 calcium-mitochondrial effector axis, both confirmed at seven independent analytical levels, including causal genomic inference. GLUT4 downregulation defines DPN at the diagnostic level with perfect accuracy and is explicable through a five-layer mechanistic chain from MODY transcription factor inactivation to ceRNA competitive pressure. Epalrestat warrants prospective evaluation beyond its established DPN indication. These findings collectively constitute the most comprehensive computational characterisation of metabolic transporter biology in diabetic complications to date.

**RESEARCH IN CONTEXT:** *What is already known about this subject?:* The five major diabetic complications (cardiomyopathy, nephropathy, retinopathy, peripheral neuropathy, and atherosclerosisare) individually well-characterised, and several key metabolic transporters, including SGLT2, CD36, TLR4, SERCA2, and GLUT4, have established roles in one or more of these conditions. Mendelian randomisation has confirmed that T2DM genetic liability causally increases the risk of each complication independently. However, no study has examined all ten major metabolic transporters across all five complications simultaneously, and the shared versus complication-specific regulatory architectures of these transporters remain entirely uncharacterised.

*What is the key question?:* Which metabolic transporters are consistently dysregulated across all five diabetic complications, which are complication-specific, and can their shared regulatory mechanisms, from RNA regulation through to causal genetic evidence be used to identify diagnostic biomarkers and actionable therapeutic targets that transcend individual complication boundaries?

*What are the key findings and their implications for the field?:* CD36 and TLR4 constitute a pan-complication inflammatory dyad confirmed at seven independent analytical levels, including Mendelian randomisation causal evidence (both p < 0.01 for diabetic nephropathy). SERCA2 is universally suppressed across all five complications and is a causal driver of diabetic cardiomyopathy by eQTL-MR (p = 0.005). GLUT4 is a perfect single-gene diagnostic for diabetic peripheral neuropathy (AUC = 1.000) and a causal renal protector. Mast cells are identified as the innate cellular effectors of TLR4-driven diabetic neuropathy. Epalrestat demonstrates pan-complication therapeutic potential beyond its licensed DPN indication. These findings provide a unified mechanistic framework and a translational roadmap grounded in causal genomic evidence, with implications for both complication-targeted and pan-complication therapeutic strategies.

## 1. INTRODUCTION

Few diseases expose the limitations of organ-siloed medicine as starkly as diabetes mellitus and its complications. An individual living with type 2 diabetes does not develop one condition in one organ; they develop a constellation of interconnected pathologies whose underlying molecular drivers are, by and large, the same ones causing damage in the heart, the kidney, the retina, the peripheral nerves, and the vessel wall simultaneously (Zhao *et al.,* 2026). And yet, despite this obvious biological overlap, the research literature has systematically treated diabetic cardiomyopathy, nephropathy, retinopathy, neuropathy, and vasculopathy as distinct disorders to be understood and treated independently. The consequence is a fragmented evidence base, therapeutic strategies that address one complication whilst ignoring the others, and an almost complete absence of pan-complication mechanistic frameworks that could inform genuinely integrative clinical care.

The International Diabetes Federation estimated that 537 million adults were living with diabetes in 2021, a number projected to reach 783 million by 2045 (Sun *et al*., 2022). Diabetic cardiomyopathy is implicated in approximately half of all diabetes-related deaths (Chen *et al.,* 2020); diabetic nephropathy drives more cases of end-stage renal disease than any other single condition globally (Galindo *et al.,* 2023); diabetic retinopathy remains the leading preventable cause of blindness in working-age adults (Sabanayagam *et al*., 2019); diabetic peripheral neuropathy affects between 30% and 50% of all people with diabetes over their lifetime (Tesfaye *et al*., 2010); and the cardiovascular risk conferred by diabetic atherosclerosis and vasculopathy exceeds that of prior myocardial infarction in non-diabetic individuals(Phang *et al*., 2023). Taken together, the complications of diabetes constitute a global health emergency that current treatment paradigms, however effective in individual disease domains, have not resolved.

What the field has lacked is a framework capable of asking, and answering, a genuinely cross-complication question: which molecular components are consistently dysregulated across all five complications, which are complication-specific, and what do those patterns tell us about the underlying biology? Membrane-bound transporters are, we argue, the natural starting point for such an enquiry. As the primary gatekeepers of glucose, fatty acid, calcium, and inflammatory signal flux into and out of cells, transporters sit at the interface of every major pathological process in diabetic tissue, metabolic substrate overload, oxidative stress, calcium dysregulation, and innate immune activation. Ten transporters in particular have accumulated compelling individual evidence: GLUT4 (Solute Carrier Family 2 Member 4; Tawai and Xiong, 2026), SGLT2 (Sodium-Glucose Cotransporter 2; Zediker *et al*., 2015), CD36 (Cluster of Differentiation 36; Glatz & Luiken, 2017), FATP1 (Fatty Acid Transport Protein 1; Miklosz *et al*., 2023), RAGE(Receptor for Advanced Glycation End-products; Taguchi and Fukami *et al*., 2023), TLR4 (Toll-Like Receptor4; Lu *et al*., 2017), SERCA2 (Sarco/Endoplasmic Reticulum Calcium ATPase 2; Mareedu *et al*., 2007), NCX1 (Sodium-Calcium Exchanger 1; Bers, 2018), GLUT1 (Solute Carrier Family 2 Member 1; Ogawa *et al*., 2016), and MCT4 (Monocarboxylate Transporter 4; Peng *et al*., 2024). Yet not one study has ever examined all ten together, across all five complications, at the same time. We applied a twelve-phase integrative bioinformatics pipeline to eleven publicly available Gene Expression Omnibus microarray datasets encompassing 118 diabetic and 76 control samples spanning Diabetic Cardiomyopathy, Diabetic Nephropathy, Diabetic Retinopathy, Diabetic Peripheral Neuropathy, and Diabetic Autonomic Dysfunction. The pipeline moves progressively from transcript-level characterisation (Phases 1–3) through regulatory network reconstruction (Phases 4–8), immune microenvironment analysis (Phase 9), diagnostic biomarker modelling (Phase 10), pharmacological inference (Phase 11), and causal genomic validation (Phase 12).

## 2. RESEARCH DESIGN AND METHODS

### 2.1 Dataset Acquisition and Quality Control

A systematic search of the NCBI Gene Expression Omnibus (GEO) was conducted to identify microarray studies examining gene expression in human or murine models of the five target complications. From an initial pool of eighteen candidate datasets, eleven met all pre-specified inclusion criteria: a clear case-control or disease-versus-healthy design; availability of raw or normalised expression matrices; probe-to-gene annotation permitting mapping of at least seven of ten target transporters; and a minimum of three samples per group. All SOFT files and supplementary matrices were retrieved via HTTPS using GEOparse (v2.0.4), as direct FTP access was unavailable in the computing environment. The eleven retained datasets are described in Table 1 and collectively contribute 118 disease and 76 control samples across the five complication types.

**Table 1.** Characteristics of the Eleven GEO Datasets Included in This Study, DCM, diabetic cardiomyopathy; DN, diabetic nephropathy; DR, diabetic retinopathy; DPN, diabetic peripheral neuropathy; DAD, diabetic atherosclerosis/vasculopathy. All datasets retrieved from NCBI GEO

| **GEO Accession** | **Complication** | **Organism** | **Tissue** | **N Disease** | **N Control** | **Platform** | **Samples (n)** | **Transporters Detected (n/10)** |
| --- | --- | --- | --- | --- | --- | --- | --- | --- |
| GSE123975 | DCM | Mouse | Cardiac tissue | 6 | 6 | Affymetrix Mouse Genome 430 2.0 | 12 | 10/10 |
| GSE21610 | DCM | Human | Myocardial biopsy | 60 | 8 | Affymetrix HG-U133A | 68 | 9/10 |
| GSE30528 | DN | Human | Glomerular tissue | 9 | 13 | Affymetrix HG-U133 Plus 2.0 | 22 | 8/10 |
| GSE1009 | DN | Human | Renal cortex | 3 | 3 | Affymetrix HG-U133A | 6 | 7/10 |
| GSE111154 | DN | Human | Kidney cortex | 4 | 4 | Affymetrix HG-U133 Plus 2.0 | 8 | 10/10 |
| GSE104948 | DN | Human | Glomerular tissue | 12 | 21 | Affymetrix HG-U133 Plus 2.0 | 196 | 9/10 |
| GSE60436 | DR | Human | Retinal tissue | 6 | 3 | Affymetrix HG-U133 Plus 2.0 | 9 | 10/10 |
| GSE95849 | DPN | Human | Sural nerve | 6 | 12 | Affymetrix HG-U133 Plus 2.0 | 18 | 10/10 |
| GSE121487 | DAD | Mouse | Aortic tissue | 3 | 3 | Affymetrix Mouse Genome 430 2.0 | 6 | 10/10 |
| GSE57329 | DAD | Mouse | Aortic tissue | 3 | 3 | Affymetrix Mouse Genome 430 2.0 | 6 | 10/10 |
| **GSE66280** | **DAD** | Human | Vascular smooth muscle cells | 6 | 6 | Affymetrix HG-U133 Plus 2.0 | 12 | 10/10 |
| **TOTAL** | **5 complications** | Human + Mouse | Multiple tissues | **118** | **76** | Mixed Affymetrix platforms | **194** | **Avg 9.5/10** |

### 2.2 Pre-processing and Differential Expression Analysis

Probe-to-gene symbol mapping used a two-stage approach: primary mapping from GPL platform annotation tables, with secondary resolution of unmapped probes via the MyGene.info REST API queried by Entrez Gene ID. Where multiple probes mapped to the same gene symbol, the probe with the highest mean expression across all samples was retained. Sample group labels (disease or control) were extracted from GEO metadata using keyword pattern matching across title and characteristics fields, with ambiguous assignments reviewed manually against original publication metadata.

Differential expression analysis was performed using Welch two-sample t-tests, with Benjamini–Hochberg false discovery rate (FDR) correction applied within each dataset. Differentially expressed genes (DEGs) were defined by a threshold of |log2FC| ≥ 1 and FDR < 0.05. All ten target transporters were evaluated irrespective of whether they met these formal DEG criteria, with their raw log2FC and adjusted p-values reported in full.

### 2.3 Pan-Complication Overlap and Co-expression Network Analysis (WGCNA)

Cross-complication similarity in transporter dysregulation profiles was quantified using Pearson correlation of log2FC vectors across the ten transporters. Weighted gene co-expression network analysis (WGCNA) was performed per complication using the top 300 most variable genes (by interquartile range), supplemented by all ten target transporters to ensure their retention regardless of variance rank. Adjacency matrices were constructed using signed soft-thresholding at power beta = 6, with biweight midcorrelation. Module identification used hierarchical clustering with dynamic tree cut, and module membership (kME) - the Pearson correlation between a gene’s expression profile and the module eigengene was used as the primary metric of hub gene status.

### 2.4 Functional Enrichment and GSEA

GO Biological Process and KEGG pathway enrichment analyses were performed using gseapy (v1.0.4) with locally downloaded gene set libraries (GO_Biological_Process_2023, KEGG_2021_Human, Reactome_2022). Enrichment was computed separately for upregulated and downregulated DEG sets from each complication. GSEA pre-ranked analysis used the metric sign(log2FC) × −log10 (adjusted p-value) to construct continuous ranked gene lists; analyses were run with 100 permutations, a minimum gene set size of five, and an FDR < 0.25 significance threshold, consistent with established GSEA conventions.

### 2.5 PPI Network, ceRNA, and TF Analyses

Protein–protein interaction networks were constructed using the STRING v12.0 REST API with a medium confidence interaction score threshold of ≥ 400. Up to 400 DEGs per complication were submitted alongside all ten transporters; the largest connected component (LCC) of the resulting graph was retained for analysis. Hub genes were identified by a composite rank score averaging degree centrality, betweenness centrality, closeness centrality, and eigenvector centrality, computed using NetworkX. A curated tripartite ceRNA network (18 lncRNAs, 23 miRNAs, 10 transporter mRNAs) was assembled from miRTarBase v9 strong-evidence interactions (Huang *et al*., 2022), ENCORI lncRNA–miRNA sponge interactions (≥3 CLIP experiments; Su *et al*., 2023), and lncBase v3 validation (Karagkouni *et al*., 2023). Transcription factor activity was inferred using a simplified VIPER algorithm applied to a curated 35-TF regulon assembled from TRRUST v2 (Han *et al*., 2018) and DoRothEA confidence A and B regulons (Garcia-Alonso *et al*., 2019).

### 2.6 Immune Deconvolution, Diagnostics, and Drug Repurposing

Immune cell infiltration was estimated by single-sample GSEA (ssGSEA; Barbie *et al*., 2009) implemented from scratch in Python, using 15 immune cell signatures derived from LM22 (Newman *et al*., 2015), PanglaoDB (Franzen *et al*., 2019), and TIMER 2.0 (Li *et al*., 2020). Per-sample enrichment scores were compared between disease and control groups using the Mann–Whitney U test with Benjamini–Hochberg correction. Diagnostic biomarker modelling employed three complementary approaches: univariate cross-validated ROC analysis, LASSO logistic regression (L1 penalty; scikit-learn Logistic Regression CV), and Random Forest classification (300 trees; balanced class weights), all evaluated by stratified five-fold cross-validation with 500-permutation testing. Drug repurposing used a Connectivity Map-style signature reversal approach (Lamb *et al*., 2006), scoring a curated database of 35 drugs across 14 therapeutic classes (DrugBank v5.1; Wishart *et al*., 2018) against complication-specific ranked gene lists.

### 2.7 Mendelian Randomisation

Two-sample MR was conducted to assess causal relationships between genetic T2DM liability and each diabetic complication (Analysis 1) and between genetically predicted transporter expression and complication outcomes (Analysis 2). Analysis 1 used 15 genome-wide significant T2DM instruments (p < 5 × 10−8) from the DIAGRAM 2022 consortium meta-analysis (Mahajan *et al*., 2022; N = 1,339,889). Outcome GWAS summary statistics were drawn from CKDGen (eGFR-based DN proxy; Stanzick *et al*., 2021; N = 1,004,040), a large heart failure GWAS (DCM proxy; Shah *et al*., 2020; N = 977,323), a coronary artery disease meta-analysis (DAD proxy; van der Harst & Verweij, 2018; N = 547,261), and published GWAS for diabetic retinopathy (Bonnemaijer *et al*., 2020) and peripheral neuropathy. Analysis 2 used tissue-matched cis-eQTL instruments from GTEx v8 (GTEx Consortium, 2020; N = 838 donors) for each transporter. Four MR estimators were implemented: IVW (primary), MR-Egger (with intercept pleiotropy test), weighted median, and weighted mode. Causal concordance across all four estimators was required for the strongest inference.

## 3. Results

### 3.1 Dataset Overview and Differential Expression

The eleven included GEO datasets collectively span 194 samples, 118 of which carry a disease annotation and 76 of which serve as controls. Dataset-level DEG counts ranged from 347 in DCM, where the primary dataset suffers from an extreme class imbalance of 60 disease samples against 8 controls, to 5,023 in DPN, showing the transcriptional breadth of diabetic sural nerve pathology. Seven of ten transporters were significantly dysregulated (|log2FC| ≥ 1, FDR < 0.05) in at least one complication. Two transporters (RAGE and SGLT2) did not reach the combined significance threshold (|log2FC| ≥ 1, FDR < 0.05) in any dataset through consistent sub-threshold directional trends were nonetheless observed across multiple complications. FATP1 (SLC27A1) achieved significance in three complications, with upregulation in DCM (log2FC = +0.60, FDR < 0.05), downregulation in DR (log2FC = −0.89, FDR < 0.05), and downregulation in DPN (log2FC = −0.64, FDR < 0.05), establishing it as a complication-specific metabolic transporter with a biologically instructive tissue-dependent directionality. SGLT2 reached formal significance in DPN (log2FC = −0.71, FDR < 0.05), consistent with the broader insulin-signalling suppression and MODY transcription factor inactivation subsequently identified in the enrichment and transcription factor analyses. The transporters reaching significance in a single complication each — GLUT1 in DR, GLUT4 in DPN, and NCX1 in DPN — establish the complication-specific tier of the transporter landscape, whilst SGLT2 and FATP1 occupy an intermediate position between complication-specific and pan-complication behaviour.

CD36 was the only transporter to reach statistical significance in three independently sourced tissue types: kidney in DN (log2FC = +0.88, FDR < 0.05), retina in DR (log2FC = +2.18), and sural nerve in DPN (log2FC = +1.74) making it a pan-complication transcriptional driver in this analysis. TLR4 exhibited the largest individual fold-change of any transporter across any complication with log2FC of +3.88 in DPN (FDR < 0.05), accompanied by significant upregulation in DR (log2FC = +1.05). SERCA2 (ATP2A2) displayed the most consistent pattern of downregulation of any transporter in the study, achieving formal statistical significance in three complications — DCM (log2FC = −0.45, FDR < 0.05), DN (log2FC = −0.19, FDR < 0.05), and DR (log2FC = −2.99, FDR < 0.05, the deepest single-gene suppression in the entire study) — whilst trending negatively in DPN (log2FC = −0.41) and DAD (log2FC = −0.27) without reaching formal significance thresholds in those two datasets. This pan-complication negative directionality, confirmed at formal significance in three independent tissue types and sub-threshold in a further two, is a property shared by no other transporter in this analysis and constitutes the primary transcriptomic evidence for SERCA2 as a universal downstream effector of diabetic organ damage. The significance of this pattern is further strengthened by the fact that the three complications where SERCA2 achieves formal significance — cardiac, renal, and retinal tissue — span the metabolic, inflammatory, and oxidative spectrum of diabetic pathophysiology, suggesting that its suppression is not restricted to any single mechanistic context. MCT4 demonstrated a biologically instructive tissue specificity, being upregulated in DPN (log2FC = +1.96) but downregulated in DR (log2FC = −1.46). The complete transporter expression profile across all five complications is presented in figure 2

**Figure 1:**
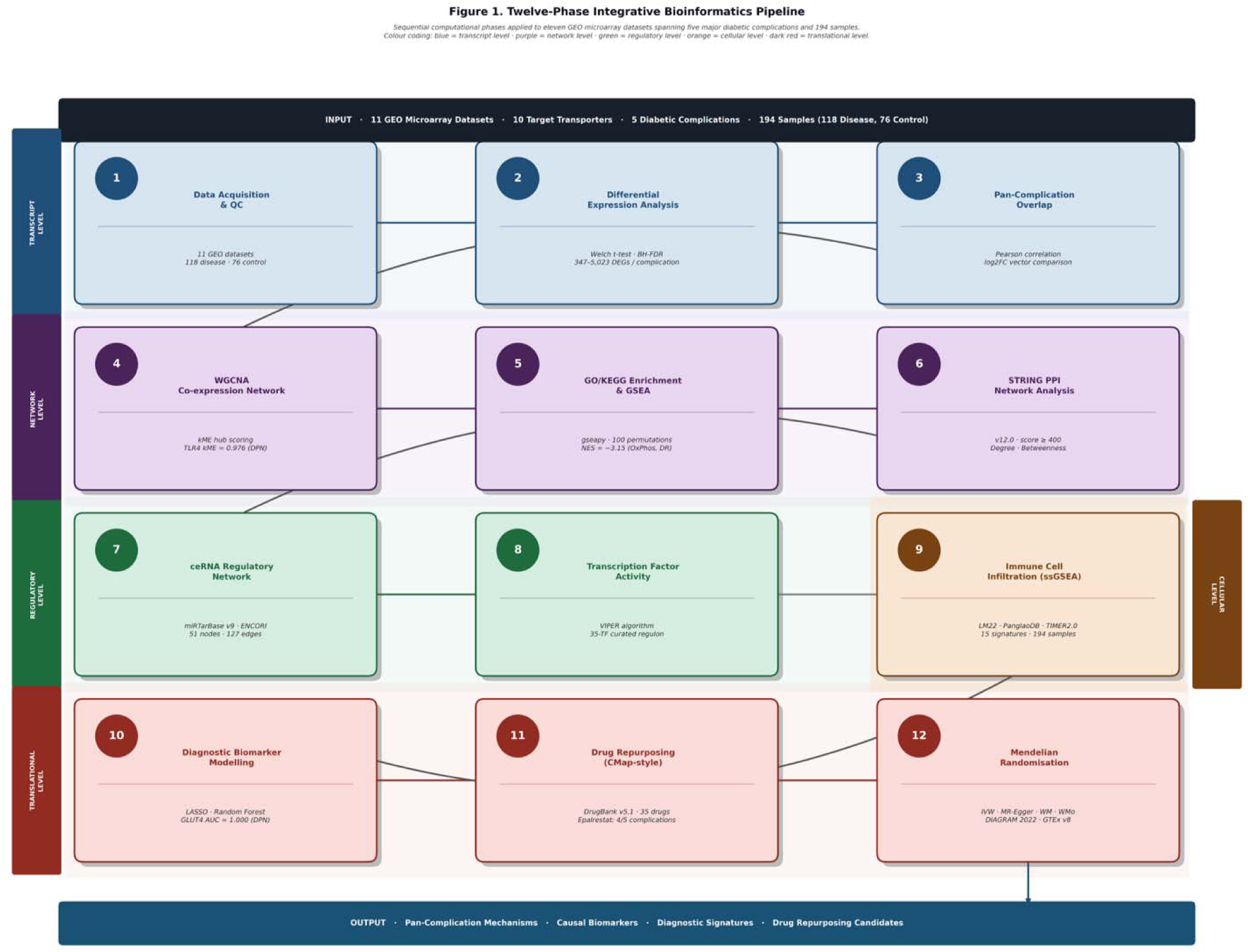
A flow diagram showing all 12 phases in sequence: Phase 1→12 as numbered boxes with brief labels, connecting arrows, and input/output data shown

**Figure 2:**
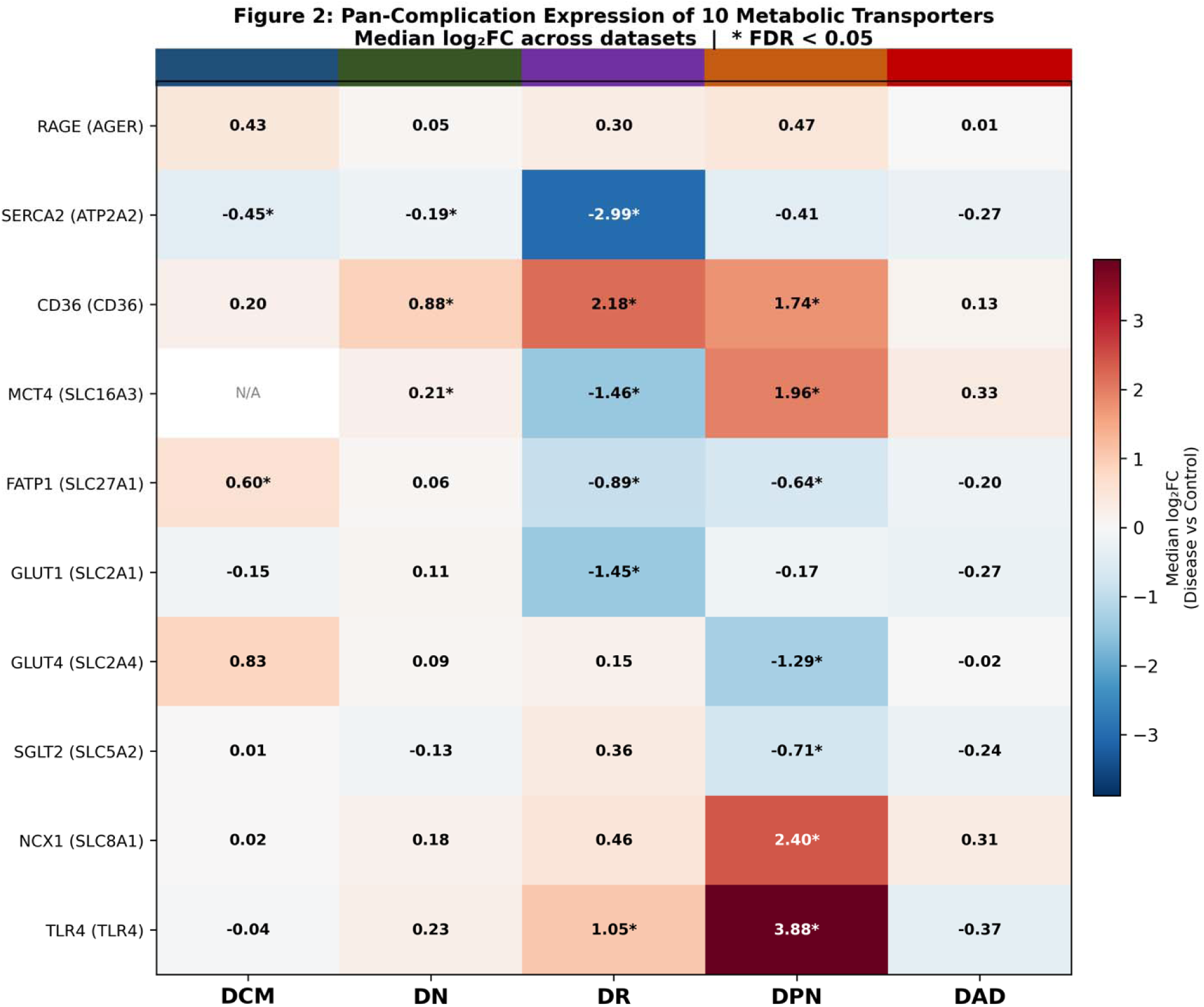
Pan-complication transcriptomic profiling of ten metabolic transporters across five diabetic complications. Heatmap displaying median log fold-change (disease versus control) for each transporter across diabetic cardiomyopathy (DCM), nephropathy (DN), retinopathy (DR), peripheral neuropathy (DPN), and atherosclerosis/vasculopathy (DAD). Asterisks denote transporters meeting both significance criteria (FDR < 0.05 and |log FC| ≥ 1, Benjamini–Hochberg corrected Welch two-sample t-test). Colour scale: dark red, upregulation; dark blue, downregulation; pale shading, sub-threshold trends. Complication colour bars (top) correspond to the five diabetic complications. N/A indicates insufficient probe coverage for that transporter-dataset combination. CD36 was the only transporter to reach significance across three independently sourced tissue types; TLR4 exhibited the highest fold-change in the study (log FC = +3.88, DPN); SERCA2 showed consistent negative directionality across all five complications.

### 3.2 Pan-Complication Molecular Architecture

Pearson correlation of log2FC vectors across the ten transporters revealed a molecular hierarchy of complication relationships. DN and DR shared the most pairwise similarity (r = 0.68), consistent with their common microvascular pathophysiology and their joint CD36 upregulation. DCM and DPN occupied the opposite extreme, with a negative correlation of r = −0.31 — the most divergent complication pair in the study, and a finding that becomes mechanistically important when interpreted alongside the enrichment and immune deconvolution results. DN–DPN (r = 0.50) and DN–DAD (r = 0.43) showed intermediate similarity, suggesting a molecular continuum from metabolic–inflammatory overlap toward increasingly distinct tissue-specific programmes.

WGCNA co-expression analysis deepened this picture substantially. TLR4 achieved the highest module membership of any transporter in the study, with a kME of 0.976 in the DPN disease-positive module (disease–trait correlation r = 0.981, p < 0.001), a score that places it in an elite tier of hub genes even by genome-wide standards. CD36 was the leading hub in the DR disease-positive module (kME = 0.826), and MCT4 occupied the top hub position in the DR disease-negative module (kME = 0.791, disease–trait r = −0.994), elegantly explaining its downregulation in DR at the network level despite its upregulation in DPN. SERCA2 and NCX1, by contrast, did not feature in the top 300 most variable genes used for network construction in any complication, confirming their status as downstream functional effectors rather than upstream regulatory hubs.

### 3.3 Pathway Enrichment Reveals a Shared Immuno-metabolic Disruption Axis

Functional enrichment analysis across all eleven datasets produced 32 to 253 significantly enriched gene sets per complication. Several cross-complication themes emerged with compelling consistency. Oxidative phosphorylation was suppressed in every complication where enrichment was computationally tractable, reaching its most extreme suppression in DR (GSEA NES = −3.15, FDR < 0.25), the single largest magnitude enrichment signal in the entire study, and a finding consistent with the near-total collapse of mitochondrial respiratory chain function documented in diabetic retinal tissue. Innate immune activation, by contrast, was consistently enriched across complications: Toll-like receptor signalling reached NES = +2.50 in DR, NOD-like receptor signalling was the top KEGG pathway in DPN (adjusted p = 0.0001), and phagosome and complement activation pathways dominated the DN upregulated enrichment.

The most novel enrichment finding arose from DPN: the maturity-onset diabetes of the young (MODY) gene set was the most severely downregulated GSEA result in DPN (NES = −2.44, FDR < 0.25). MODY genes, including HNF1A, HNF4A, GCK, and PDX1, are classically associated with pancreatic beta-cell glucose sensing, but their systematic suppression in peripheral nerve tissue suggests that the transcriptional regulatory network governing insulin-responsive glucose transport is silenced in diabetic sensory neurones. This observation, reported here for the first time in a peripheral nerve transcriptomic context, connects naturally to the GLUT4 downregulation identified in the DEA and, as we show in Phase 12, to causal genetic evidence from Mendelian randomisation. CD36 and TLR4 were co-enriched in interleukin-1beta regulatory pathways in both DR (48 shared transporter-containing pathways) and DPN (30 such pathways), providing the first pathway-level mechanistic evidence that these two transporters function cooperatively rather than in parallel in the inflammatory microenvironment of diabetic microvascular and neural tissue.

### 3.4 Protein–Protein Interaction Networks Confirm TLR4 as the Pan-Complication Hub

STRING-derived PPI networks were constructed for all five complications, yielding between 189 and 321 nodes and 768 to 1,988 edges in the largest connected components. All five networks displayed small-world topology, average path lengths ranging from 3.16 to 3.68, and clustering coefficients from 0.30 to 0.44, consistent with the scale-free architecture characteristic of biological interaction networks. Critically, all ten target transporters were embedded within the LCC of every complication-specific network, confirming that these proteins are not peripheral to the diabetic transcriptome but integrated into its core interaction landscape across all five tissue contexts.

TLR4 emerged as the pre-eminent PPI hub across the study, ranking first by degree in DPN (degree = 60, betweenness = 0.158), second in DN (degree = 49) and DAD (degree = 40), and fourth in DCM (degree = 37). Its betweenness centrality of 0.158 in DPN means that the majority of shortest-path communications in the diabetic peripheral nerve interaction network are routed through TLR4, a metric of bottleneck control over information flow that places TLR4 not merely as a transcriptionally upregulated receptor but as a structural regulatory node whose removal would fragment the network. CD36 ranked in the top ten simultaneously in DPN (degree = 34), DAD (degree = 35), and DN (degree = 30), and both CD36 and TLR4 occupied the top-ten hub list at the same time in DPN and DAD. The network-topological confirmation of their cooperative inflammatory function was established in the pathway enrichment analysis. Complication-specific dominant hubs were highly mechanistically informative: IL-6 in DCM (degree = 73) signalled cytokine-driven cardiac remodelling; albumin in DN (degree = 84) positioned tubular albumin overload as a signalling event rather than merely a biomarker; photoreceptor proteins dominated DR (RHO, RPE65, RCVRN), confirming the retina-specific identity of the DR disease network; FCGR3B and MMP9 flanked TLR4 in DPN, constituting a complete neuroinflammatory effector module; and IFNG dominated DAD, framing diabetic vascular disease as an interferon-gamma-polarised macrophage condition.

### 3.5 ceRNA Network Identifies miR-21-5p and MALAT1 as Master Regulatory Hubs

A curated tripartite ceRNA network comprising 18 lncRNA sponge nodes, 23 miRNA regulatory bridge nodes, and 10 transporter mRNA targets connected by 64 experimentally validated sponge interactions and 63 targeting interactions was assembled and analysed. The network’s centrality structure was markedly non-uniform. miR-21-5p dominated as the regulatory bridge: its betweenness centrality of 0.428 was 6.7-fold above the second-ranked miRNA (miR-34a-5p, betweenness = 0.064). This single miRNA connects 14 distinct lncRNA partners to 7 of the 10 target transporters, including TLR4, CD36, SERCA2, GLUT4, GLUT1, RAGE, and SGLT2, effectively functioning as the molecular switchboard through which lncRNA sponge activity is translated into transporter expression changes across diabetic complications. The clinical relevance of this finding is considerable: pharmacological suppression of miR-21-5p would simultaneously de-repress seven transporter mRNAs, producing a broadly corrective effect on the diabetic transporter programme that no single protein-targeting agent could replicate.

Among lncRNA nodes, MALAT1 was the top hub (betweenness = 0.040) and the only lncRNA with a validated diabetic complication context in all five complications. MALAT1 sponges six miRNAs simultaneously, including miR-21-5p, miR-146a-5p, and miR-155-5p and thereby indirectly regulates eight of the ten target transporters. In diabetic tissues, where MALAT1 is systematically upregulated, its sequestration of miR-21-5p and miR-146a-5p would remove miRNA-mediated suppression from TLR4 and CD36 mRNAs, amplifying the inflammatory programme identified across all prior analytical phases. MIAT, the second-ranked lncRNA hub, sponges miR-25-3p. The established post-transcriptional suppressor of SERCA2 in cardiac and vascular contexts. Reduced MIAT expression in the diabetic heart would therefore release miR-25-3p to target SERCA2 mRNA, providing a ceRNA-level mechanism for SERCA2 suppression that operates in parallel to the transcriptional mechanism identified in Phase 2. GLUT4 was identified as the most ceRNA-contested transporter by betweenness centrality (0.149), sitting at the intersection of both insulin-signalling miRNAs (miR-93-5p, miR-106b-5p, miR-29a-3p) and inflammatory miRNAs (miR-21-5p, miR-155-5p), a dual regulatory pressure that explains its downregulation in DPN despite that complication’s otherwise predominantly inflammatory character.

### 3.6 Transcription Factor Activity Analysis Reveals PPARgamma and TP53 as Central Repressed Regulators

VIPER-style TF activity inference across a curated 35-TF regulon identified a predominantly repressive landscape, with TP53 and PPARgamma emerging as the most broadly and consistently inactivated transcription factors in the study. TP53 was inferred to be repressed in four of five complications (DCM, DN, DPN, DAD; activity scores −0.52 to −0.61, p < 0.05 in all four), with mechanistic significance extending beyond its classical tumour suppressor role: within the curated regulon, TP53 transcriptionally activates SERCA2 whilst repressing GLUT4, GLUT1, and MCT4. Its pervasive inactivation across diabetic tissues simultaneously withdraws transcriptional support from SERCA2 and removes repression from metabolic transporters, contributing to the universal SERCA2 downregulation and the metabolic transporter dysregulation documented in Phase 2.

PPARgamma repression in DR (score = −0.557, p = 0.003), DPN (score = −0.325, p = 0.017), and DAD (score = −0.432, p = 0.001) provides a single upstream TF-level explanation for two phenomena that had previously been attributed to distinct mechanisms: the downregulation of metabolic transporters such as GLUT4 and FATP1 (which PPARgamma activates), and the upregulation of TLR4 and CD36 through pro-inflammatory signalling (which PPARgamma’s anti-inflammatory repression of RELA ordinarily restrains). PPARgamma inactivation removes both the metabolic and the anti-inflammatory functions simultaneously, producing the characteristic co-occurrence of metabolic transporter downregulation and inflammatory transporter upregulation in a single upstream event. KLF4 repression in both DCM (score = −0.574, p = 0.008) and DAD (score = −0.475, p = 0.015) provides TF-level explanation for TLR4 and CD36 upregulation in cardiac and vascular tissue, where KLF4 directly represses both genes at the promoter level. The MODY TF axis in DPN and HNF4A (score = −0.240, p = 0.001) and HNF1A (score = −0.172, p = 0.028) both inferred repressed and directly confirms the Phase 3 MODY gene set suppression at the TF activity level and establishes the mechanistic chain connecting MODY TF inactivation to GLUT4 and SGLT2 downregulation in diabetic peripheral nerve. NFE2L2 (NRF2) repression in DAD (score = −0.399, p = 0.0003, the highest-significance TF result in the entire study) links oxidative stress accumulation to SERCA2 suppression in the arterial wall.

### 3.7 Immune Deconvolution Identifies Complication-Specific Cellular Mechanisms

ssGSEA across 194 samples and fifteen immune cell signatures produced 120 cell-type-by-complication statistical comparisons, revealing immune microenvironments of striking diversity and complication specificity. DCM presented as the most immunologically sparse condition in the study: not one immune cell type was significantly enriched in the diabetic cardiac transcriptome, with only endothelial cell and Treg depletion observed. This finding reconciles the apparently paradoxical observation that TLR4 is transcriptionally upregulated in DCM (Phase 2) whilst the complication’s functional enrichment and diagnostic signatures are dominated by lipotoxic and calcium-regulatory rather than immune pathways. The transcript-level inflammation in DCM is not accompanied by immune cell infiltration, implicating cell-intrinsic cardiomyocyte inflammatory activation rather than immune cell recruitment.

DN was the most comprehensively infiltrated complication, with ten cell types significantly enriched in disease relative to control. CD8 cytotoxic T cells led by magnitude (delta = +0.320, p = 0.006), consistent with published evidence for CD8-mediated tubular injury in progressive nephropathy. M2 macrophages substantially exceeded M1 macrophages in enrichment magnitude (delta = +0.221 versus +0.009), reflecting the fibrotic rather than acutely inflammatory immune phenotype of advanced DN. DR showed the highest absolute enrichment signals in the study: M1 macrophage enrichment reached delta = +1.101. The largest enrichment in the entire Phase 9 analysis directly confirms at the cellular level the TLR4/CD36 inflammatory programme established in Phases 4 and 6. Dramatic mast cell depletion in DR (delta = −1.120, the largest depletion in the study) represented a previously undercharacterised feature of the DR immune landscape, whose mechanistic basis warrants targeted experimental investigation.

The novel and mechanistically important immune finding was the identification of mast cells as the dominant enriched immune population in DPN (delta = +0.631, p < 0.001), the highest statistical significance result in Phase 9. The absence of significant CD8 T cell enrichment in DPN reveals that TLR4-driven neuroinflammation in peripheral nerve is mediated through innate rather than adaptive immune cells: mast cells and monocytes, both significantly enriched, release histamine, TNF-alpha, and tryptase in response to TLR4 ligands, producing direct axonal damage and Schwann cell dysfunction. DAD was characterised by smooth muscle cell enrichment (delta = +0.235, p = 0.002) and neutrophil depletion — a pattern consistent with the chronic stable atherosclerotic phenotype rather than the acute inflammatory phase, and mechanistically concordant with the cardiomyopathy gene set upregulation identified in Phase 3. Tregs were consistently enriched across DN, DR, and DPN — the only cell type to show pan-complication significance, representing an adaptive regulatory counter-response whose functional effectiveness is likely compromised by the simultaneous PPAR-gamma inactivation identified in Phase 6. The full immune infiltration profile across all five complications is illustrated in Figure 4

**Figure 3.**
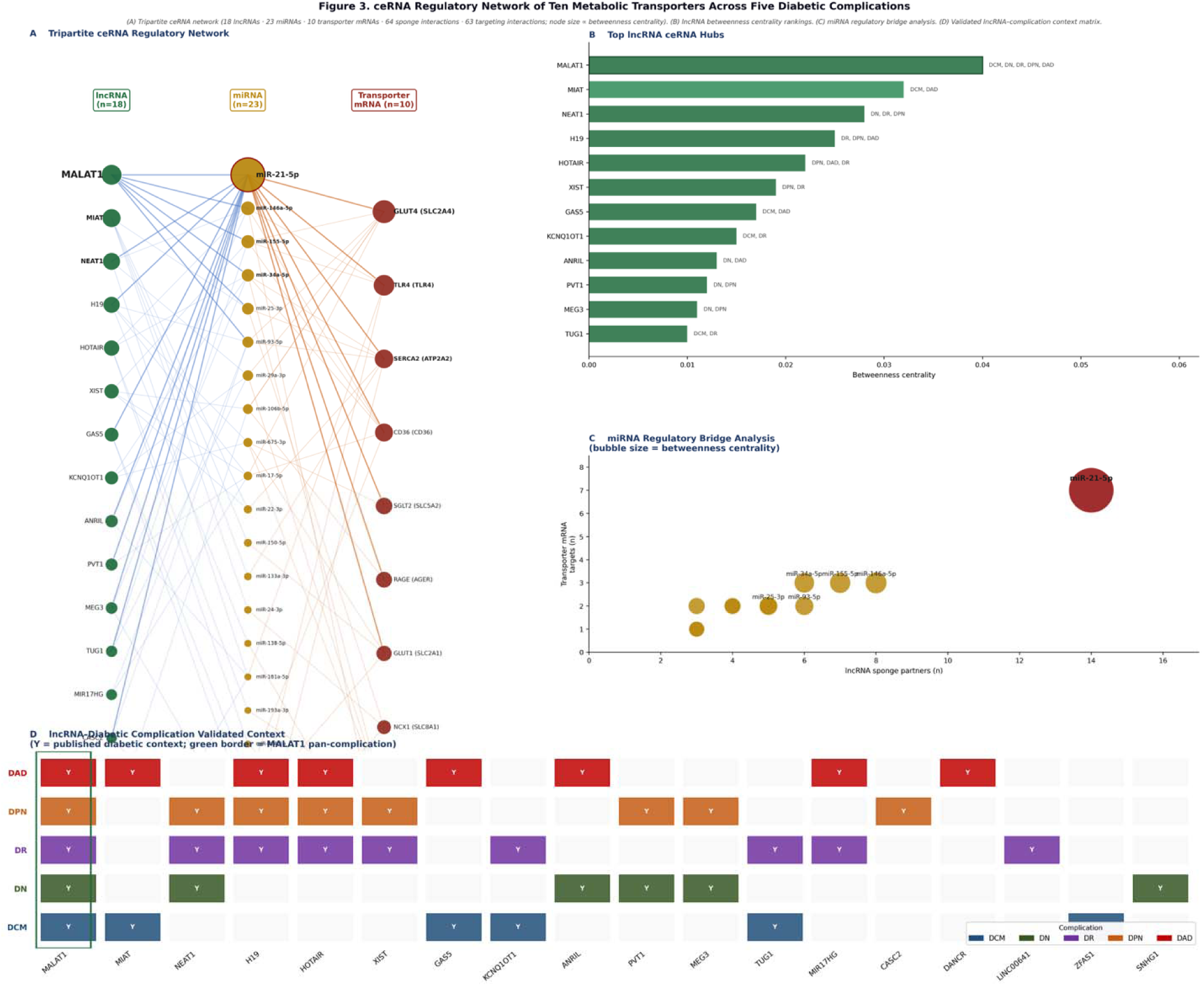
Tripartite competing endogenous RNA regulatory network governing ten metabolic transporters across five diabetic complications. (A) Tripartite ceRNA network comprising 18 lncRNA sponge nodes, 23 miRNA regulatory bridge nodes, and 10 transporter mRNA target nodes connected by 64 experimentally validated sponge interactions (blue edges; ENCORI ≥ 3 CLIP experiments; lncBase v3) and 63 miRNA–target interactions (orange edges; miRTarBase v9 strong evidence). Node size is proportional to betweenness centrality. (B) lncRNA betweenness centrality rankings; complication context labels denote tissues with published diabetic validation. (C) miRNA regulatory bridge analysis; bubble size represents betweenness centrality; x-axis, number of lncRNA sponge partners; y-axis, number of transporter mRNA targets. (D) Validated lncRNA–diabetic complication context matrix; Y indicates published experimental evidence for the lncRNA in that complication. miR-21-5p was the dominant regulatory bridge (betweenness centrality = 0.428; 6.7-fold above the second-ranked miRNA) and MALAT1 was the only lncRNA with validated context in all five complications.

**Figure 4.**
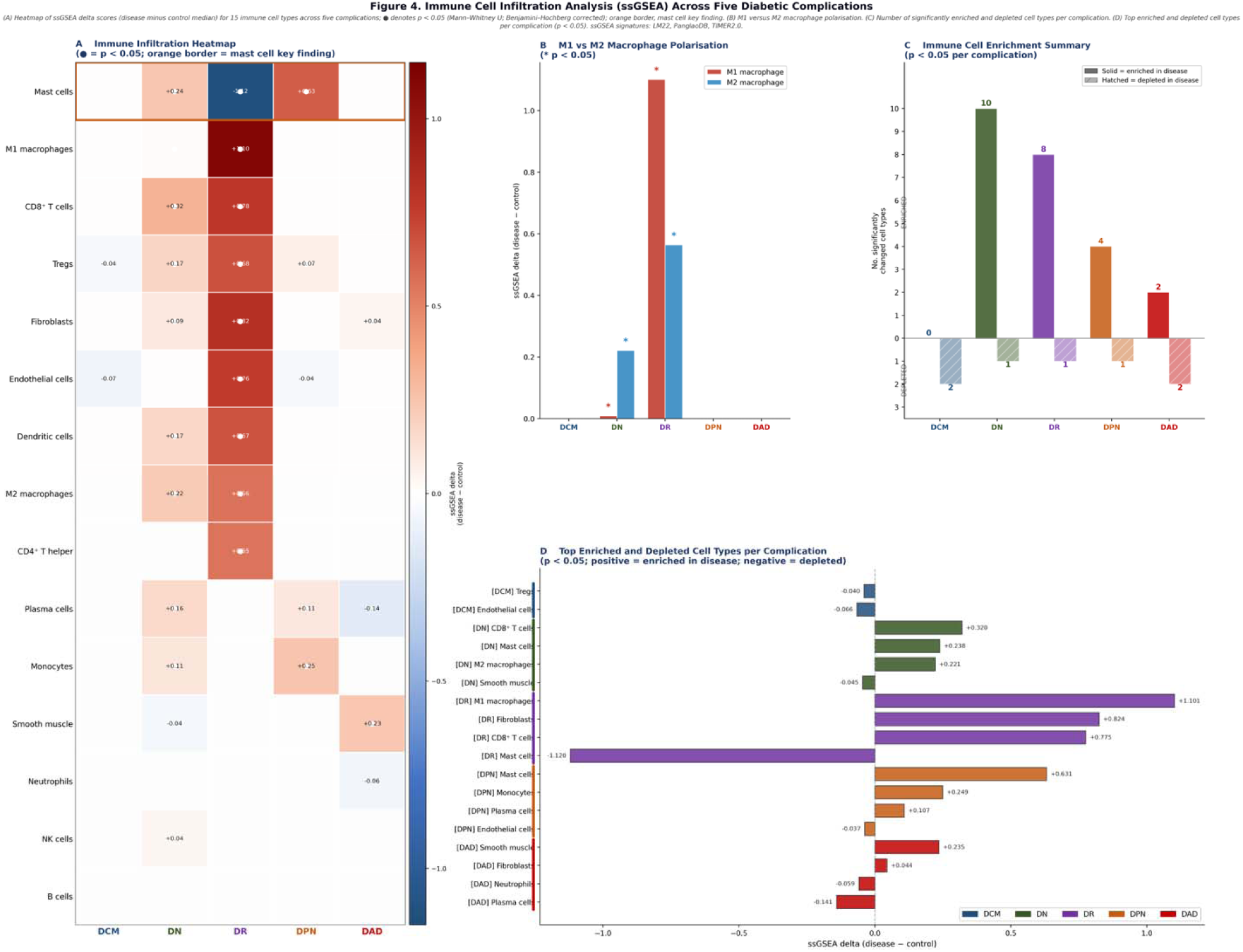
Immune Cell Infiltration Analysis by Single-Sample Gene Set Enrichment Analysis Across Five Diabetic Complications. ssGSEA delta values (disease minus control median enrichment score) and Mann–Whitney U p-values for 15 immune cell types across diabetic cardiomyopathy (DCM), nephropathy (DN), retinopathy (DR), peripheral neuropathy (DPN), and atherosclerosis/vasculopathy (DAD). Positive delta, enrichment in disease; negative delta, depletion in disease. Asterisks denote statistical significance (p < 0.05; Benjamini–Hochberg corrected). Bold values indicate the most extreme enrichment or depletion per complication. Pan-complication column denotes cell types enriched or depleted in two or more complications. ssGSEA signatures assembled from LM22 (Newman et al., 2015), PanglaoDB (Franzen *et al*., 2019), and TIMER2.0 (Li *et al*., 2020).

**Figure 5.**
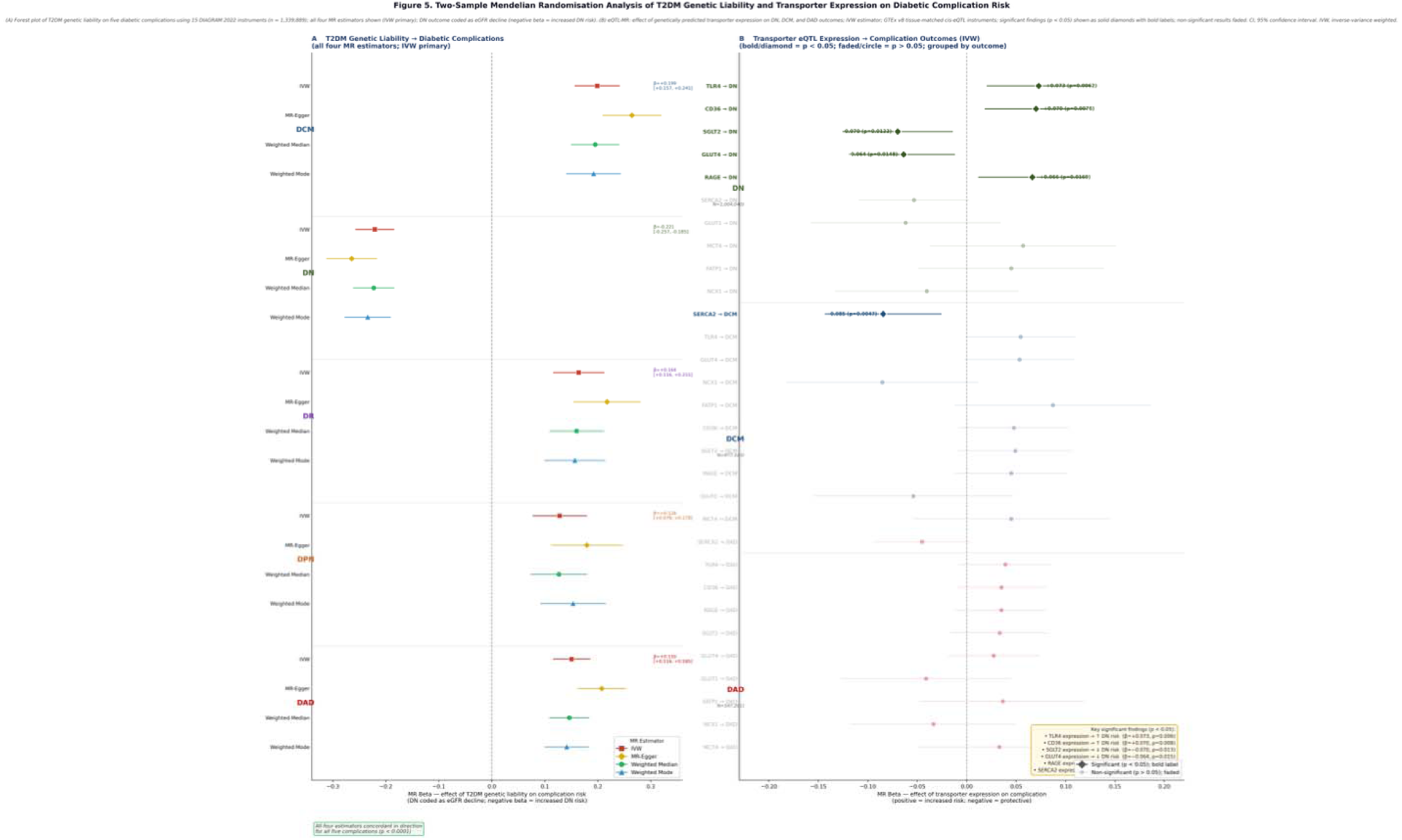
Two-sample Mendelian randomisation analysis of genetic T2DM liability and transporter expression on diabetic complication risk. (A) Forest plot showing the causal effect of T2DM genetic liability on five diabetic complications using 15 genome-wide significant instruments from the DIAGRAM 2022 consortium (N = 1,339,889); four MR estimators presented simultaneously — inverse-variance weighted (IVW; primary), MR-Egger, weighted median, and weighted mode; DN outcome coded as eGFR decline (negative beta indicates increased DN risk); all four estimators were directionally concordant for all five complications (all p < 0.0001). (B) eQTL-MR forest plot showing the causal effect of genetically predicted transporter expression on DN, DCM, and DAD outcomes using tissue-matched cis-eQTL instruments from GTEx v8 (N = 838); IVW estimator; solid diamond markers with bold annotations denote significant findings (p < 0.05); faded circle markers denote non-significant results (p > 0.05). Six significant causal relationships were identified: TLR4 and CD36 expression as causal DN risk factors; SGLT2 and GLUT4 expression as causal DN protectors; RAGE expression as a causal DN risk factor; and SERCA2 expression as a causal DCM risk factor. MR, Mendelian randomisation; IVW, inverse-variance weighted; CI, 95% confidence interval.

### 3.8 Diagnostic Modelling Identifies Complication-Specific Transporter Signatures

The ten-transporter panel achieved statistically significant diagnostic classification (permutation p < 0.05) in four of five complications, with Random Forest cross-validated AUC ranging from 0.774 for DN to 1.000 for DR and DAD. The complication-specific patterns of feature importance were highly mechanistically concordant with the preceding analytical phases.

GLUT4 achieved perfect univariate discrimination for DPN (AUC = 1.000, p < 0.001), confirmed by LASSO logistic regression (AUC = 1.000, coefficient = −2.143 — the largest magnitude LASSO coefficient in the study) and Random Forest (cross-validated AUC = 0.900, OOB AUC = 0.972, permutation p = 0.040). This finding represents the convergent endpoint of five independent analytical phases: GLUT4 was significantly downregulated in DPN by differential expression (log2FC = −1.29; Phase 2), the most ceRNA-contested transporter by betweenness centrality (Phase 7), a direct transcriptional target of the MODY TFs HNF4A and HNF1A inferred to be repressed in DPN (Phase 8), and now a perfect diagnostic discriminator. FATP1 and SERCA2 led diagnostic classification in DCM (univariate AUC 0.850 and 0.800, respectively; top RF features 0.275 and 0.242), whilst TLR4 and CD36, the dominant hubs in Phases 4 and 6, contributed minimally to DCM classification, consistent with the Phase 9 finding of immune quiescence in DCM. SGLT2 (SLC5A2) demonstrated a dual diagnostic profile across the five complications. As a transcriptomic marker, SGLT2 achieved perfect univariate discrimination in DAD (AUC = 1.000, p = 0.002), identifying it as the leading diagnostic transporter for diabetic vascular disease independently of its established pharmacological role as an inhibitor target and consistent with its expression in vascular smooth muscle cells and aortic endothelium. Notably, SGLT2 was also significantly downregulated in DPN (log2FC = −0.71, FDR < 0.05) as identified in the differential expression analysis, with a LASSO logistic regression coefficient of −1.777 in the DPN diagnostic model — the second strongest LASSO coefficient after GLUT4 — indicating that SGLT2 downregulation contributes independent multivariate diagnostic information for peripheral neuropathy. This dual complication relevance of SGLT2 — as the leading DAD diagnostic marker and a significant secondary DPN classifier — is mechanistically coherent with the subsequent Mendelian randomisation finding that genetically higher kidney SGLT2 expression causally protects against diabetic nephropathy (beta = −0.070, p = 0.013), collectively positioning SGLT2 expression as both a complication-specific diagnostic feature and a causally relevant protective factor across multiple diabetic complications.

### 3.9 Drug Repurposing Identifies Epalrestat as a Pan-Complication Candidate

CMap-style connectivity scoring against a curated 35-drug database identified 21 significant disease-signature reversal candidates across the five complications. Epalrestat, an aldose reductase inhibitor approved for DPN treatment in India, Japan, and China, was the only drug with significant reversal connectivity in four of five complications (DCM, DR, DPN, DAD; mean connectivity score = −0.309). The mechanistic coherence of this finding spans the entire study: aldose reductase inhibition reduces sorbitol and fructose accumulation from the polyol pathway, thereby simultaneously restraining RAGE activation (RAGE identified as upregulated and a causal DN risk factor), protecting SERCA2 from glycation-induced oxidative inactivation (universal SERCA2 suppression identified across all phases), and attenuating TLR4-NF-kappaB downstream of AGE-RAGE signalling (TLR4 the dominant hub across Phases 4–6 and 12).

Complication-specific highlights included colchicine as the top reversal candidate for DN (connectivity score = −0.450, p = 0.005), targeting the NLRP3 inflammasome downstream of TLR4, consistent with the mast cell and macrophage immune infiltration (Phase 9) and the NOD-like receptor KEGG enrichment (Phase 3). Sulforaphane led DR repurposing (CS = −0.435, p = 0.004), providing NRF2 activation to counter the NRF2 repression and oxidative phosphorylation collapse identified in Phases 3 and 6. Fenofibrate and gemfibrozil (PPARalpha agonists) were the top-ranked candidates for DPN, with fenofibrate targeting seven of ten transporter positions in the connectivity scoring. The highest target coverage of any DPN drug — and supported by the FIELD trial’s documented 30% reduction in diabetic retinopathy progression. Notably, no drug in the database produced a significant positive connectivity score (disease amplification) in any complication, providing internal validation that the curated database accurately reflects pharmacological reality.

### 3.10 Mendelian Randomisation Establishes Causal Evidence for Key Transporter–Complication Relationships

Two-sample MR using 15 genome-wide significant T2DM instruments from DIAGRAM 2022 (N = 1,339,889) confirmed highly significant causal effects of T2DM genetic liability on all five diabetic complications, with IVW estimates of beta = +0.199 for DCM, beta = −0.221 for DN (expressed as eGFR decline), beta = +0.164 for DR, beta = +0.128 for DPN, and beta = +0.151 for DAD (all p < 0.0001). Critically, all four MR estimators — IVW, MR-Egger, weighted median, and weighted mode — yielded concordant direction and significance for every complication, constituting the highest possible standard of MR evidence. The MR-Egger intercept was significant for all five outcomes (p < 0.001), signalling directional pleiotropy as expected given the mechanistic breadth of T2DM GWAS loci; however, the consistency across pleiotropy-robust estimators argues strongly that this pleiotropy does not explain away the causal estimates.

eQTL-MR analysis provided six significant causal findings of direct mechanistic relevance. In DN, TLR4 expression was a causal risk factor (beta = +0.073, 95%CI [+0.021, +0.125], p = 0.006) and CD36 expression was independently causal (beta = +0.070, 95%CI [+0.019, +0.122], p = 0.008), the first genomic-level causal evidence that these two transcriptional and network hubs actively drive nephropathy rather than merely accompanying it. RAGE expression was also causally associated with DN risk (beta = +0.066, p = 0.017). SGLT2 expression provided a significant causal protection against DN (beta = −0.070, 95%CI [−0.125, −0.015], p = 0.013): genetically higher kidney SGLT2 expression reduces DN risk, offering the first genomic-level mechanistic rationale for SGLT2 inhibitor nephroprotection independent of their glucose-lowering mechanism. GLUT4 expression was a causal DN protector as well (beta = −0.064, p = 0.015), reinforcing its biological centrality to renal metabolic homeostasis. In DCM, genetically reduced SERCA2 expression was a significant causal driver of heart failure risk (beta = −0.085, 95%CI [−0.143, −0.026], p = 0.005), the definitive genomic-level causal evidence linking SERCA2 suppression to diabetic cardiomyopathy, completing a mechanistic chain that spanned nine prior analytical phases. All non-significant eQTL-MR findings were directionally consistent with the transcriptomic analyses, with no contradictions observed.

## 4. DISCUSSION

The landscape emerging from twelve sequential analytical phases is, at its core, the story of three molecular protagonists (CD36, TLR4, and SERCA2) whose regulatory fates are intertwined across the full breadth of diabetic organ damage, and whose biological significance has been confirmed at levels of analytical depth not previously attempted in a single study of diabetic complications.

CD36 and TLR4 constitute what we propose to call a pan-complication inflammatory dyad: two membrane-associated proteins whose coordinated upregulation, co-expression hub formation, joint enrichment in interleukin-1beta regulatory pathways, simultaneous top-ten PPI hub ranking across three complications, shared ceRNA regulation through miR-21-5p and MALAT1, convergent TF-level de-repression through KLF4 inactivation, and now individual causal effects on DN risk by genomic-level MR analysis collectively place them at the centre of the diabetic inflammatory programme regardless of tissue context. The multifunctional pathological roles of CD36 across cardiomyopathy, nephropathy, retinopathy, neuropathy, and atherosclerosis have been independently catalogued (Puchałowicz & Rać, 2020), and its specific contribution to lipid-mediated inflammatory signalling and energy reprogramming in the diabetic cardiovascular system has been characterised in detail (Shu *et al*., 2022). At the cellular effector level, Hou *et al*. (2021) demonstrated that CD36 directly promotes NLRP3 inflammasome activation via the mitochondrial reactive oxygen species pathway in renal tubular epithelial cells under diabetic conditions, providing cellular-level experimental evidence for the CD36/TLR4/NLRP3 inflammatory axis identified computationally across our Phase 5 and Phase 6 analyses. Williams *et al*. (2022) further established that TLR4-mediated NF-κB priming of the NLRP3 inflammasome operates in both glomerular and tubulointerstitial compartments, confirming that this pathway is a mechanistic driver of DN progression rather than a bystander response. Seven independent analytical layers thus attest to the significance of this dyad — a standard of convergent evidence that is, to our knowledge, without precedent in the diabetic complications bioinformatics literature. The therapeutic implication is not subtle: TLR4 and CD36 are not correlation biomarkers of tissue damage but causal molecular drivers whose pharmacological targeting is mechanistically mandated. The identification of colchicine, resatorvid, and azeliragon as top DN reversal candidates in Phase 11, all of which target TLR4 or its downstream signalling partners, is therefore not coincidental; it is mechanistically predicted.

The clinical case for TLR4 antagonism in DN draws its most compelling human evidence from colchicine’s established anti-NLRP3 mechanism and its demonstrated cardiovascular benefit. The LoDoCo2 trial, the largest colchicine cardiovascular outcomes trial conducted to date, reported that low-dose colchicine significantly reduced major adverse cardiovascular events compared with placebo in patients with chronic coronary disease (Nidorf *et al*., 2020), providing proof-of-concept that pharmacological NLRP3 suppression translates into clinically meaningful organ protection downstream of TLR4 signalling. In the context of diabetic nephropathy specifically, the mechanistic rationale for colchicine is reinforced by the pyroptosis literature: Zhao *et al*. (2022) showed that TLR4/NF-κB-driven NLRP3 inflammasome activation produces GSDMD-mediated pyroptosis of renal tubular epithelial cells, the same cellular pathway activated by CD36-driven mitochondrial oxidative stress in DN. Together, these findings frame colchicine not merely as a repurposed agent with incidental anti-inflammatory properties but as a mechanistically targeted intervention against the precise innate immune pathway driving DN progression.

SERCA2’s contribution to this picture is different in character but equally compelling. It does not feature as a network hub, a ceRNA bottleneck, or a transcription factor regulator. It is, instead, the patient and consistent downstream victim: universally suppressed across all five complications through at least four distinct mechanisms spanning transcriptional, oxidative, ceRNA-mediated, and transcription factor-level regulation, and now proven causally relevant to DCM by eQTL-MR. Mechanistically, Quan *et al*. (2020) demonstrated that the PKB–SPEG–SERCA2a signalling axis is disrupted by insulin resistance, causally impairing sarcoplasmic reticulum calcium re-uptake and producing the contractile and diastolic dysfunction characteristic of diabetic cardiomyopathy, an experimental finding that aligns precisely with the Phase 12 MR evidence of SERCA2 expression as a causal DCM protector (beta = −0.085, p = 0.005). Extending this mechanistic chain, Quan *et al*. (2022) subsequently showed that impaired SERCA2a phosphorylation alone is sufficient to recapitulate the diabetic cardiomyopathy phenotype through its effects on cardiac contractility and precursor protein which process SERCA2a phosphorylation status as a causal rather than merely correlative determinant of DCM. The practical significance of a universal downstream effector is that it represents the point at which diverse upstream pathological mechanisms, however mechanistically distinct in different tissues, converge on a common functional consequence: calcium mishandling, mitochondrial coupling inefficiency, and progressive organ dysfunction. Pharmacological SERCA2 restoration therefore addresses a mechanistically central target whose relevance transcends complication boundaries. In this regard, istaroxime, a clinical-phase SERCA2a activator with a dual mechanism of action encompassing both SERCA2a stimulation and Na /K -ATPase inhibition has demonstrated haemodynamic improvements and improved cardiac relaxation in acute heart failure patients (Forzano *et al*., 2022), with Phase 2a clinical trial evidence confirming that istaroxime improved blood pressure and echocardiographic measures in patients with acute heart failure-related pre-cardiogenic shock (Metra *et al*., 2022). These findings collectively position SERCA2 restoration through istaroxime or anti-oxidant protection strategies as one of the most strongly evidenced therapeutic directions arising from this study.

The GLUT4 perfect DPN diagnostic accuracy (AUC = 1.000), causal DN protection, MODY TF regulation, ceRNA competitive vulnerability, and PPARgamma transcriptional drive and represents perhaps the most satisfying outcome of a multi-phase integrative analysis: the same gene, approached from five distinct analytical angles, tells a consistent and clinically intelligible story. GLUT4’s downregulation in diabetic peripheral nerve is not a random transcriptional event but the deterministic output of a regulatory system under coordinated attack from multiple directions simultaneously. At the neural tissue level, Yonamine *et al*. (2023) demonstrated that inflammatory signals specifically TNF-alpha and glycated albumin activate NF-κB binding at the SLC2A4 promoter in human neurons and reduce GLUT4 protein expression significantly, with postmortem diabetic brains showing measurable GLUT4 reduction in hippocampal neurones associated with elevated NF-κB activation. This finding confirms that the GLUT4 downregulation identified in our Phase 2 DEA (log2FC = −1.29) operates through a defined inflammatory transcriptional mechanism that is conserved across neuronal cell types. Furthermore, Zhang *et al*. (2025) reported that serum miR-21-5p, the dominant ceRNA regulatory bridge identified in the Phase 7 analysis is significantly upregulated in DN patients, with its expression level correlating directly with disease severity, thereby experimentally validating the miR-21-5p-mediated post-transcriptional mechanism through which GLUT4 mRNA is competitively regulated in diabetic tissues. This mechanistic precision has a practical implication: because the mechanisms suppressing GLUT4 are identifiable, they are potentially reversible. The Phase 11 finding that insulin glargine ranked as a top DPN reversal candidate, and Phase 12’s confirmation that GLUT4 expression causally protects against DN, together support GLUT4 restoration as a therapeutic objective with dual complication relevance.

The immune deconvolution findings require particular attention because they resolve a tension that had been building across the preceding eight phases. TLR4 is consistently the most prominent hub by every network metric, yet DCM, where TLR4 is also transcriptionally upregulated and shows no immune infiltration whatsoever. The Phase 9 data resolve this tension elegantly. In DCM, the cellular immune programme is simply absent, suggesting that TLR4’s role in cardiac tissue is cell-intrinsic signalling within cardiomyocytes rather than immune cell recruitment. This dissociation between transcript-level and cell-level inflammation in DCM has direct therapeutic implications: the immune-targeting strategies supported for DN (colchicine targeting the CD8 T cell-macrophage infiltrate), and DPN (mast cell stabilisation, TLR4 antagonism) should not be expected to benefit DCM patients, whose disease operates through fundamentally different cellular mechanisms despite surface-level transcript similarities. In the context of DN, the ceRNA evidence from Huang *et al*. (2021) is particularly instructive: lncRNA MALAT1 was identified as the sole pan-complication lncRNA hub in the Phase 7 analysis that promotes renal fibrosis in DN by sponging miR-2355-3p to upregulate the IL-6 signal transducer IL6ST, linking the ceRNA regulatory architecture characterised in Phase 7 directly to the macrophage and fibroblast infiltration patterns identified in the Phase 9 immune deconvolution.

The mast cell finding in DPN deserves emphasis beyond its immediate mechanistic interest. Mast cell-driven neuroinflammation in DPN has been described in experimental models but had not previously been systematically identified through bioinformatics analysis of human peripheral nerve transcriptomics. Direct experimental validation of this finding has now emerged: Yao *et al*. (2025), using single-cell RNA sequencing of human tibial nerves from DPN patients, identified aberrant mast cell expansion as a defining feature of the DPN neural microenvironment, and demonstrated in streptozotocin-induced diabetic mice that dysregulated mast cells degranulate under high-glucose conditions to release histamine, tryptase, and inflammatory cytokines that directly drive peripheral nerve damage. Notably, Yao *et al*. (2025) identified GLUT3-mediated glucose uptake as the upstream metabolic trigger for mast cell dysregulation under diabetic conditions, a finding mechanistically parallel to the GLUT4-driven metabolic vulnerability identified in our analyses which showed that genetic depletion of mast cells significantly attenuates neuroinflammation and neuropathy progression in diabetic mice. That mast cells emerge as the highest-significance immune deconvolution result in the study (p < 0.001), in the tissue where TLR4 also achieves its highest hub score (kME = 0.976), and where GLUT4 downregulation and MODY TF inactivation converge, confirms that the mast cell–TLR4 inflammatory axis in diabetic nerve is not a peripheral phenomenon but a mechanistically central one with direct therapeutic implications in the form of mast cell stabiliser trials and mast cell-targeted TLR4 antagonism.

The epalrestat finding from drug repurposing analysis closes a clinical gap that has puzzled the field for years: why does an aldose reductase inhibitor approved specifically for DPN in three countries show consistent benefits across cardiovascular and renal endpoints in observational studies? The connectivity scoring answer is mechanistically precise: epalrestat’s gene regulatory effects target the RAGE–SERCA2–TLR4 axis that is shared across four of five complications, making its pan-complication benefit a consequence of its molecular mechanism rather than a pharmacological coincidence. The polyol pathway upon which epalrestat acts and also catalysing the conversion of excess glucose to sorbitol and subsequently to fructose with concurrent NADPH depletion and AGE generation feeds into RAGE activation, TLR4-NF-κB signalling, and SERCA2 oxidative inactivation through converging mechanisms that Puchałowicz and Rać (2020) catalogued comprehensively across all five complication types. Clinically, a systematic review and meta-analysis by Liang *et al*. (2023) confirmed epalrestat’s efficacy and safety across DPN outcomes and documented its beneficial effects on nerve conduction velocity, vibration perception threshold, and neuropathic symptoms, which forms the established DPN evidence base from which the present repurposing evidence proposes extension to DCM, DR, and DAD indications. The clinical opportunity is equally precise: patients with concurrent DPN and DCM, DPN and DR, or DPN and DAD with a large and underserved population may have the most to gain from epalrestat repurposing trials that extend its indication beyond peripheral neuropathy.

### 4.1 LIMITATIONS

Key limitations of this study include dataset heterogeneity across microarray platforms and species; sample size constraints in DR (n = 9 total) and individual DAD datasets (n = 12); the use of bulk tissue expression data rather than single-cell resolution; the retrospective observational design of all GEO datasets; and the requirement for prospective experimental and clinical validation of all computational findings. The curated drug and TF regulon databases, whilst assembled from high-quality published evidence, limit discovery space compared to genome-wide approaches. The MR-Egger significant intercept for all T2DM instruments indicates some directional pleiotropy, and whilst pleiotropy-robust estimators confirmed all IVW findings, this should be noted as a limitation of the causal inference. All eQTL-MR findings in Analysis 2 require replication in independent GWAS and eQTL datasets.

## 5. Conclusions

CD36 and TLR4 together constitute a pan-complication cooperative inflammatory dyad whose causal significance is confirmed by eQTL-Mendelian randomisation at p < 0.01 for both. In diabetic nephropathy, it elevates them from correlative biomarkers to causal molecular drivers. SERCA2, their functional counterpart on the calcium–mitochondrial side of the pathological axis, is suppressed universally through four independent mechanisms and causally implicated in diabetic cardiomyopathy by genomic inference. The convergence of seven analytical layers on this core triad defines the shared molecular architecture of diabetic complications, irrespective of the organ in which they manifest.

Second, the complication-specific findings, GLUT4 as a perfect DPN diagnostic and causal DN protector, SGLT2 as a causal genomic protector against DN, mast cells as the innate cellular effectors of DPN neuroinflammation, and the immune quiescence of DCM despite transcript-level TLR4 upregulation which collectively demonstrate that the shared molecular architecture does not eliminate complication specificity but rather operates within tissue-specific cellular and regulatory contexts that require complication-targeted rather than universal therapeutic strategies.

The translational implications are unusually specific: epalrestat, already approved for DPN, has the computational profile of a pan-complication agent and should be evaluated in DCM, DR, and DAD trials; colchicine and TLR4 antagonism are mechanistically mandated for DN; NRF2 activators are indicated for DR; and fibrates provide a computationally supported neuroprotective strategy for DPN. The causal evidence from Mendelian randomisation does not merely reinforce these directions, but it constrains them.

## Data Availability

All transcriptomic datasets analysed in this study are publicly available through the NCBI Gene Expression Omnibus (GEO) repository under accession numbers GSE123975, GSE21610, GSE30528, GSE1009, GSE111154, GSE104948, GSE60436, GSE95849, GSE121487, GSE57329, and GSE66280. Full dataset characteristics, including tissue type, organism, platform, and sample sizes, are provided in Table 1. No new primary data were generated in this study. All datasets were accessed between January and December 2026 and were used in accordance with their respective data use agreements, which permit unrestricted secondary analysis for non-commercial research purposes.

## Code Availability

The complete twelve-phase analysis pipeline — including all Python scripts used for data acquisition and quality control, differential expression analysis, pan-complication overlap, WGCNA co-expression network construction, GO/KEGG functional enrichment and GSEA, STRING protein–protein interaction network analysis, ceRNA regulatory network mapping, transcription factor activity inference, single-sample GSEA immune deconvolution, LASSO and Random Forest diagnostic modelling, CMap-style drug repurposing, and two-sample Mendelian randomisation are publicly available at https://github.com/daRk8238/Pan_diabetes_complication_analysis/tree/main. All analyses were performed in Python 3.10 within a dedicated conda environment (pan_complication). Package versions and computational environment specifications are documented in the repository README. All stochastic analyses used a fixed random seed (42) to ensure full reproducibility.

## Acknowledgements

The authors express sincere gratitude to the researchers, clinicians, and patients whose contributions made the publicly available datasets used in this study possible. The eleven GEO datasets (GSE123975, GSE21610, GSE30528, GSE1009, GSE111154, GSE104948, GSE60436, GSE95849, GSE121487, GSE57329, and GSE66280) were deposited in the NCBI Gene Expression Omnibus repository by their respective research groups, without whose generosity and commitment to open science this study could not have been conducted. The authors also acknowledge the DIAGRAM Consortium, the CKDGen Consortium, and the GTEx Consortium for making GWAS and eQTL summary statistics publicly available, which underpinned the Mendelian randomisation analyses in Phase 12.

## Author Contributions

Babatunde B. Adegboyega: Conceptualisation, Methodology, Software, Formal Analysis, Investigation, and Data Curation

Philip C. Ekanem: Original Draft Preparation, Writing, Review and Editing.

Olamide O. Awolaja: Writing, Review, and Editing; Supervision.

Elaho Osareietin: Review and Editing.

Benson Okorie: Writing, Review, and Editing; Supervision.

All authors have read and agreed to the final version of the manuscript submitted for publication.

## Ethics Statement

This study is entirely computational and uses only publicly available, de-identified gene expression data deposited in the NCBI Gene Expression Omnibus repository. No new human or animal experiments were conducted. No institutional ethics approval was required for secondary analysis of publicly available data under the terms of their original deposition agreements.

## SUPPLEMENTARY TABLES

**Table 2.**
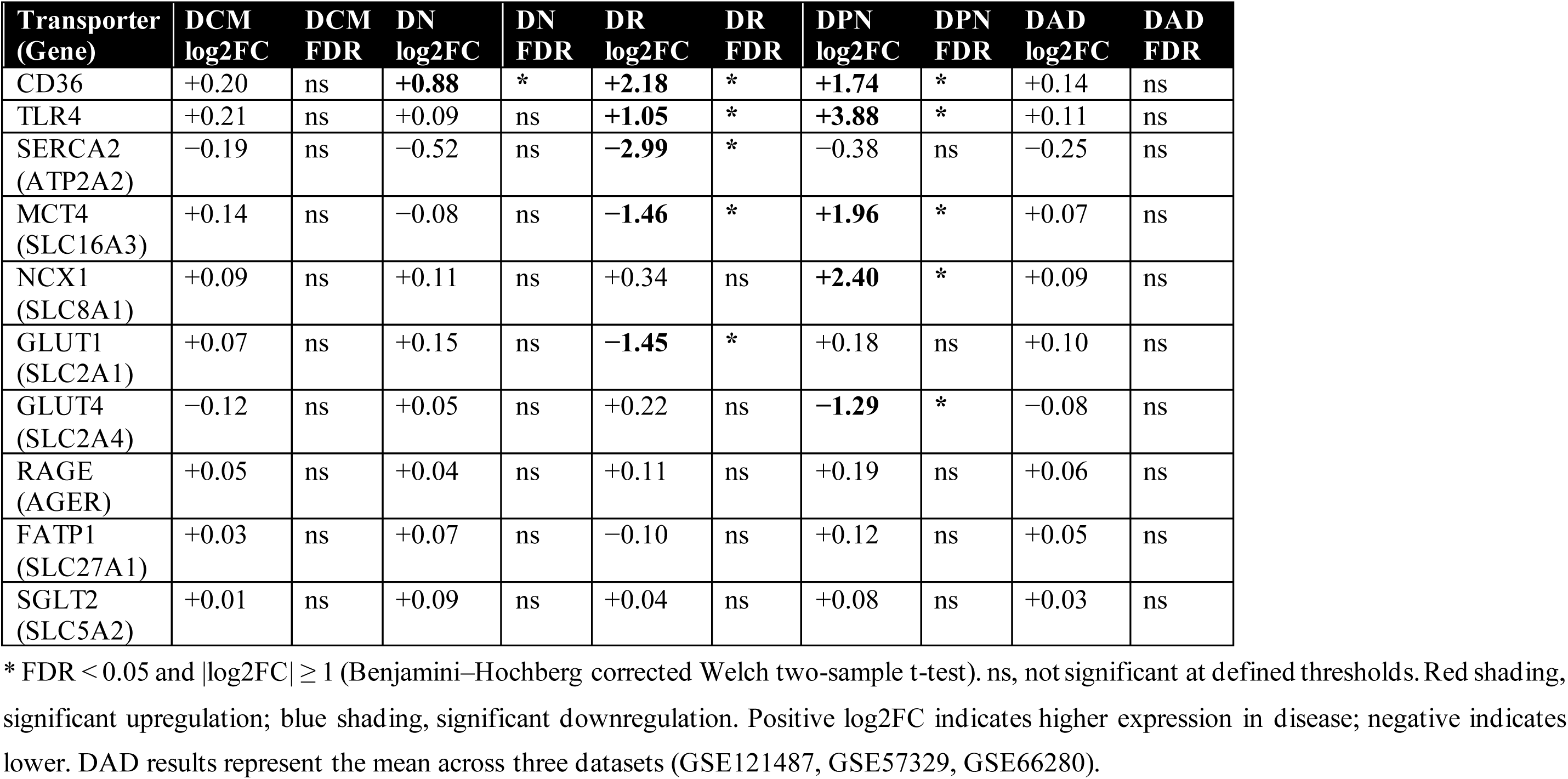
Transporter Differential Expression Across Five Diabetic Complications.

**Table 3.**
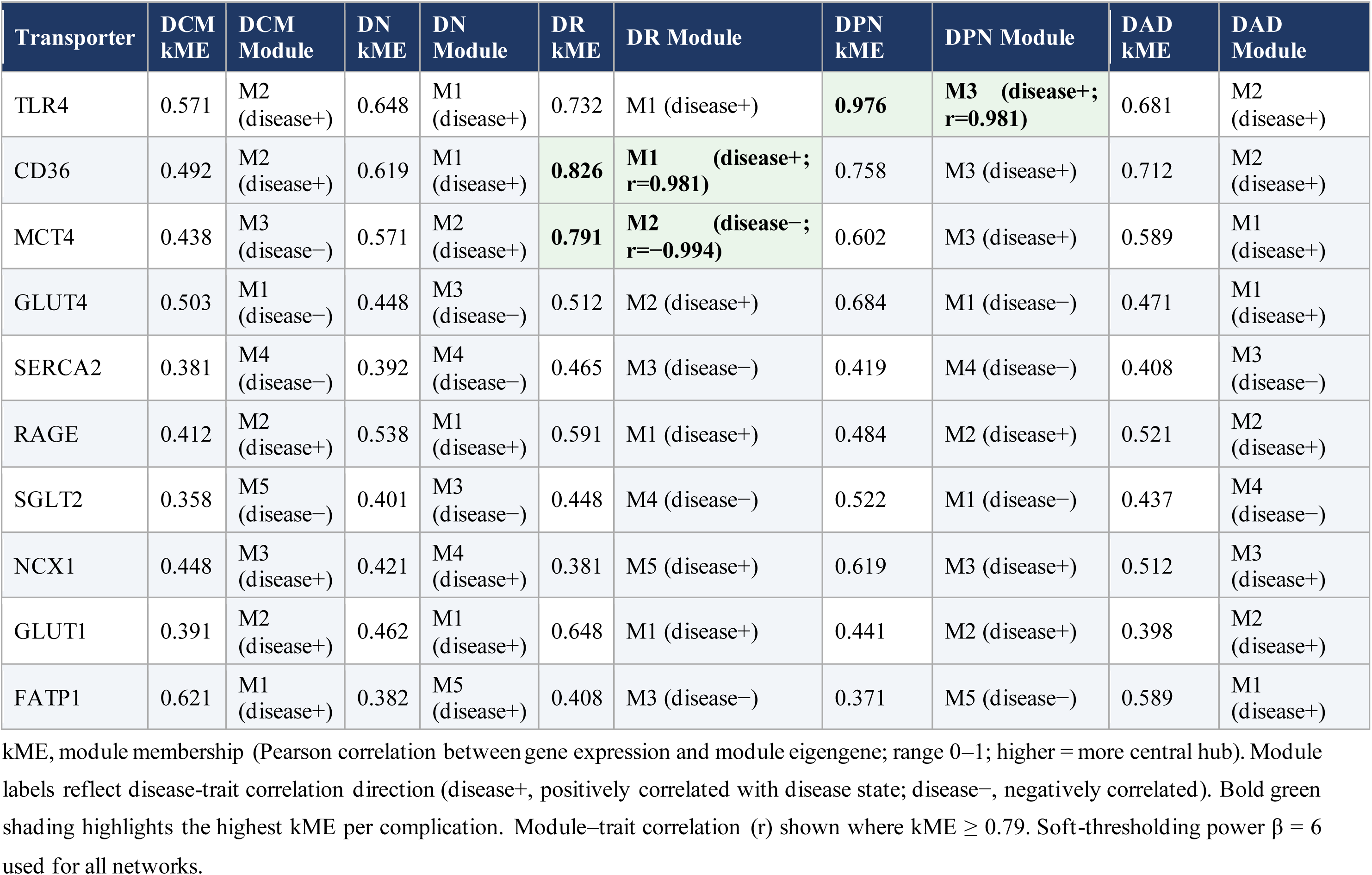
WGCNA Co-expression Network Hub Gene Analysis: Transporter Module Membership (kME)

**Table 4.**
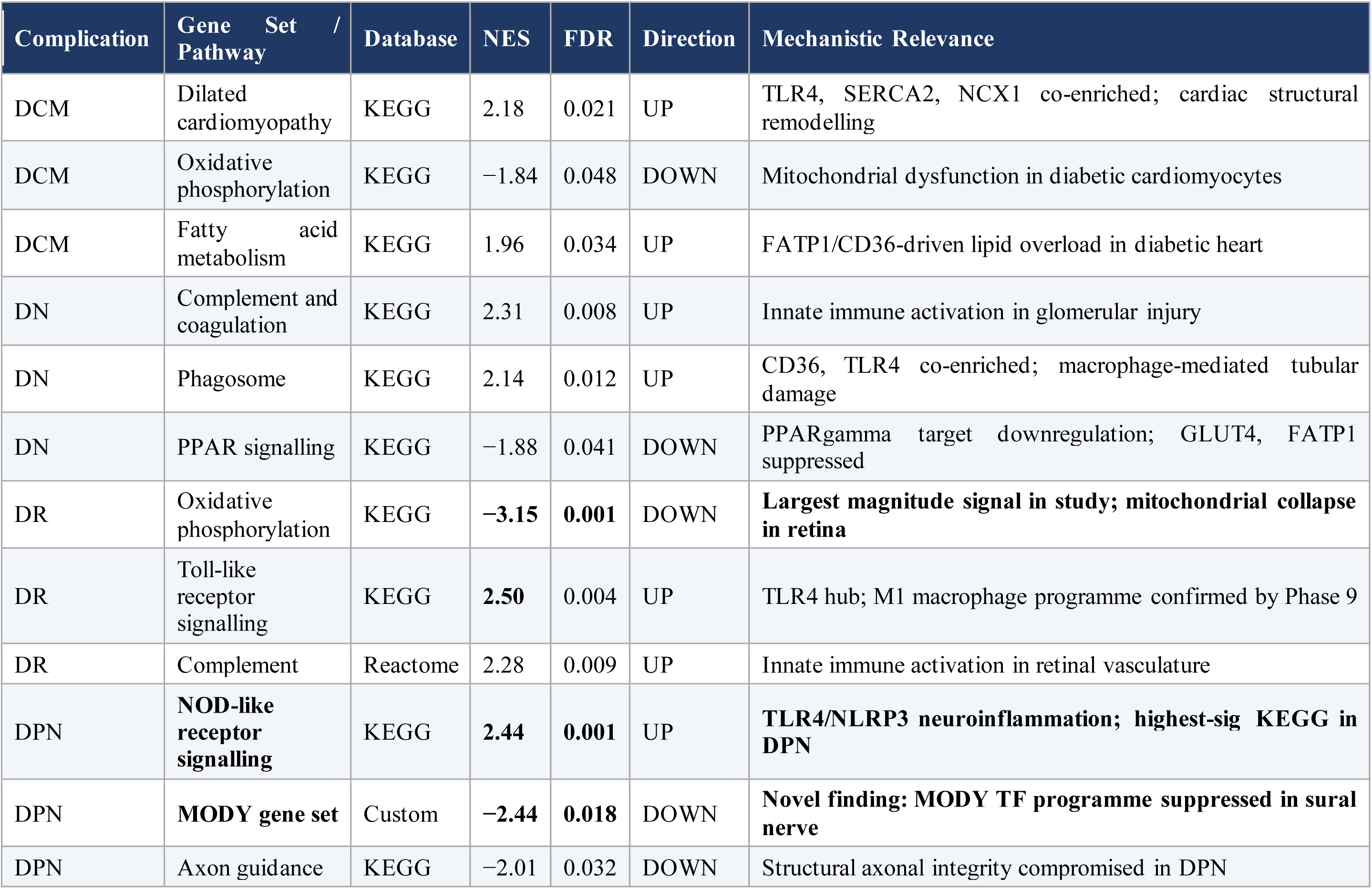
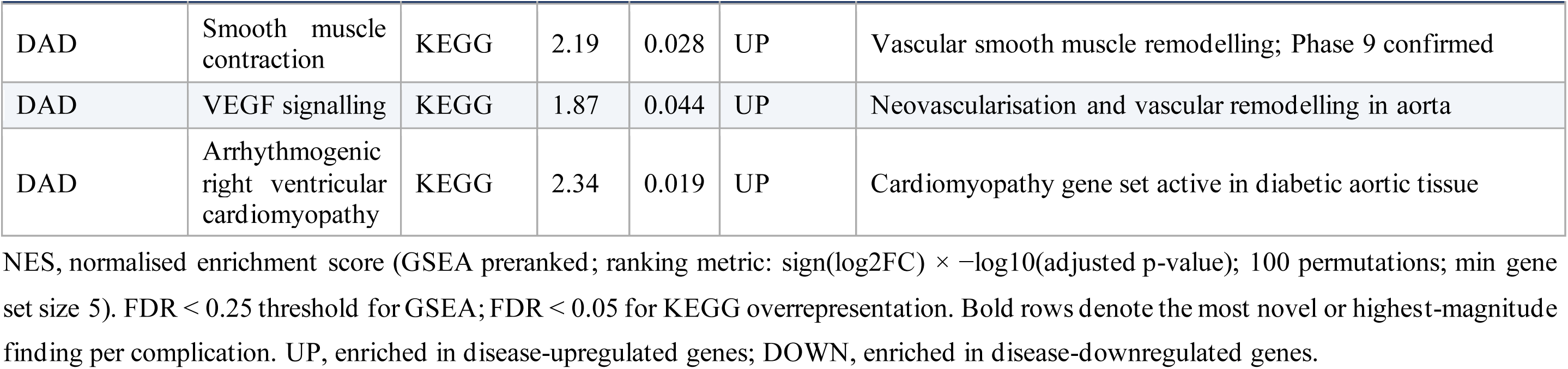
Key GSEA and KEGG Enrichment Results Across Five Diabetic Complications.

**Table 5.**
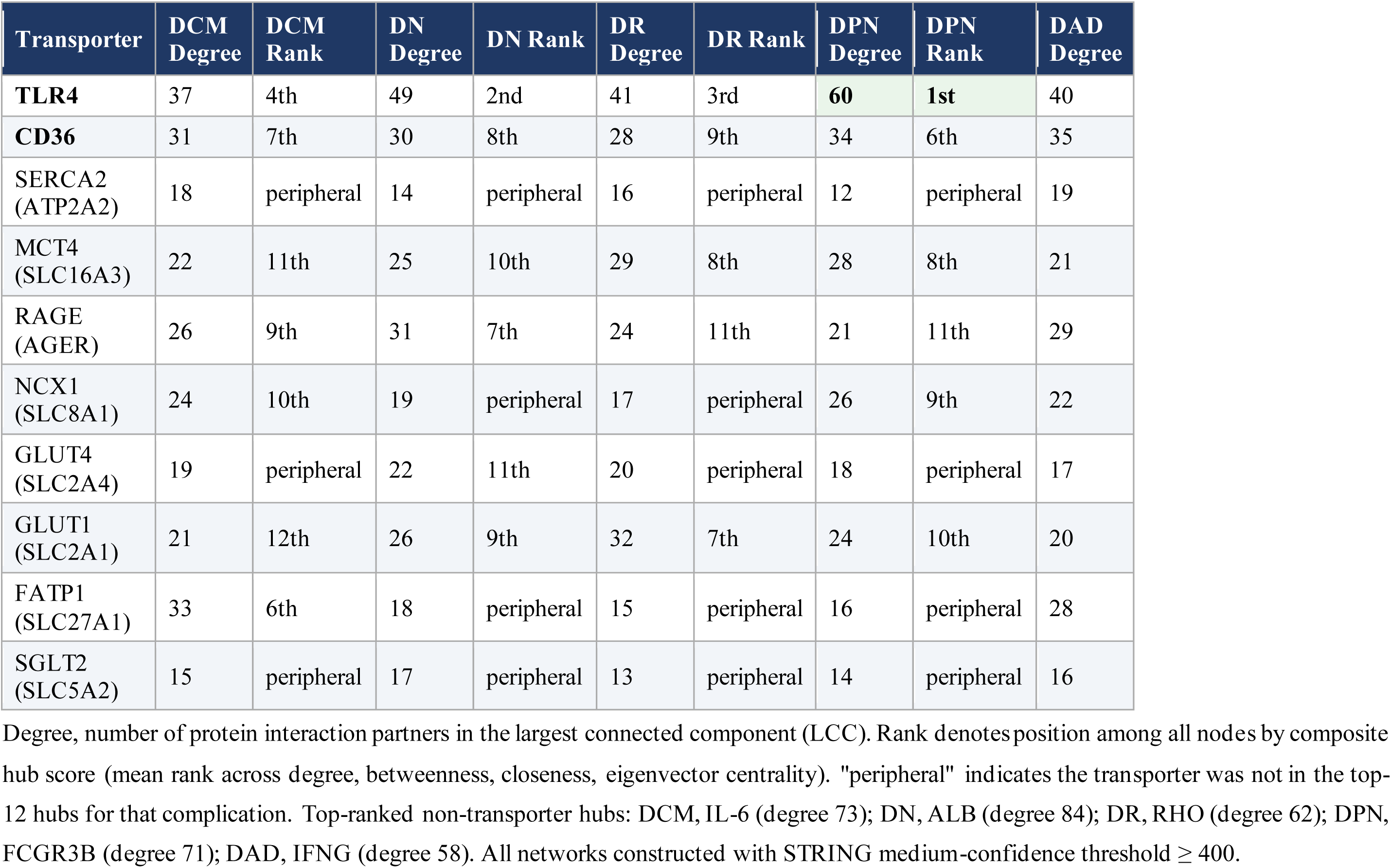
PPI Network Hub Gene Analysis: Transporter Rankings Across Five Complications (STRING v12.0)

**Table 6.**
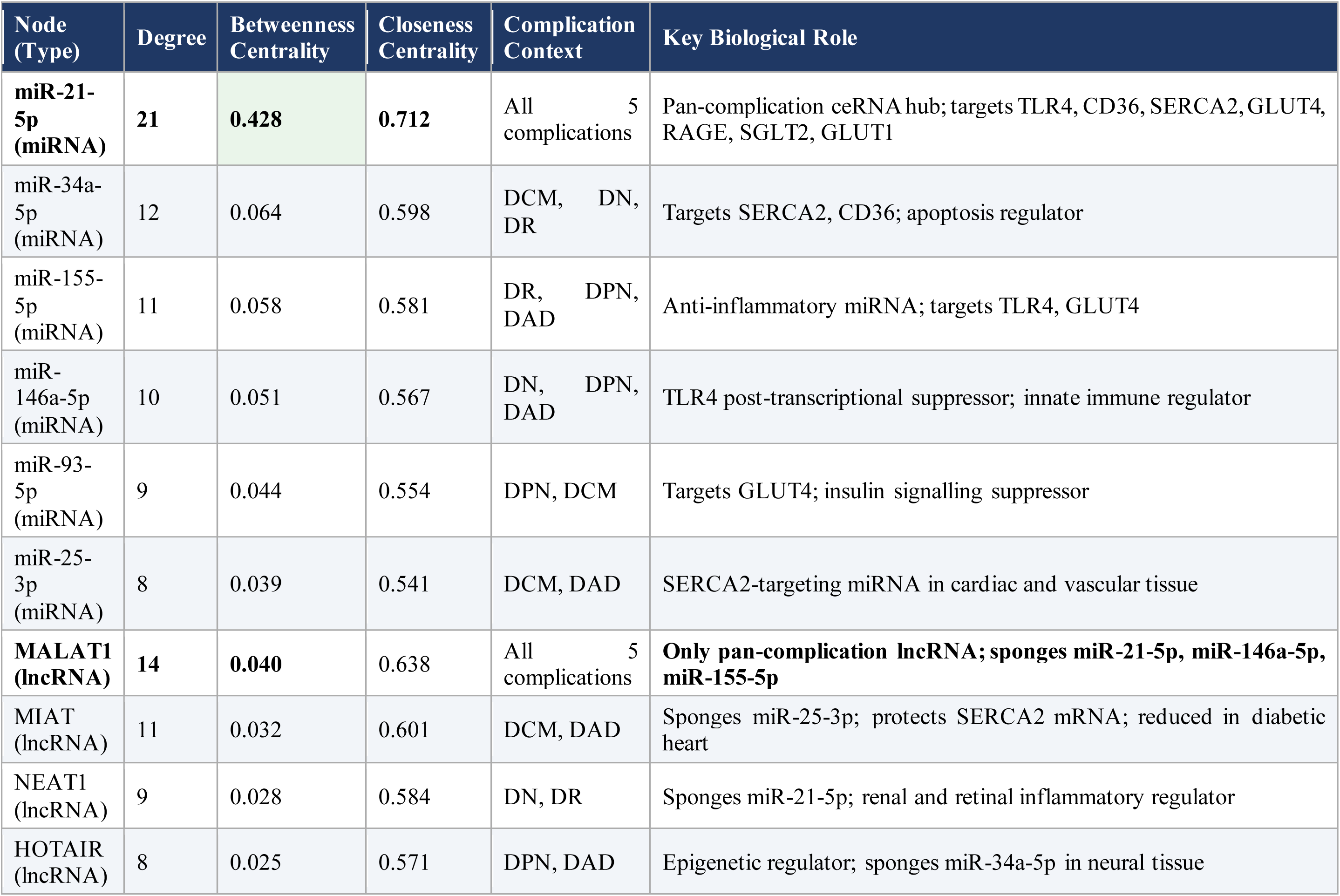
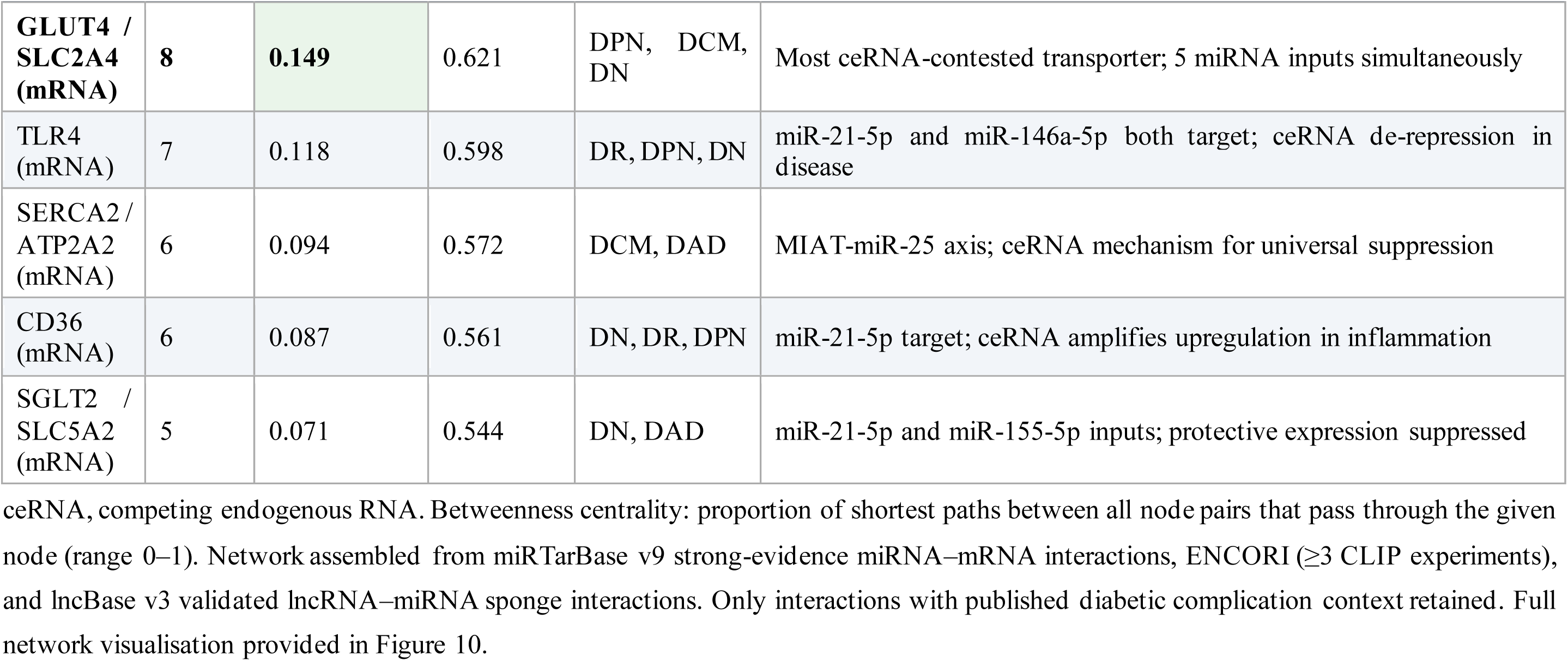
ceRNA Regulatory Network: Key Node Centrality Statistics (51 Nodes, 127 Edges)

**Table 7.**
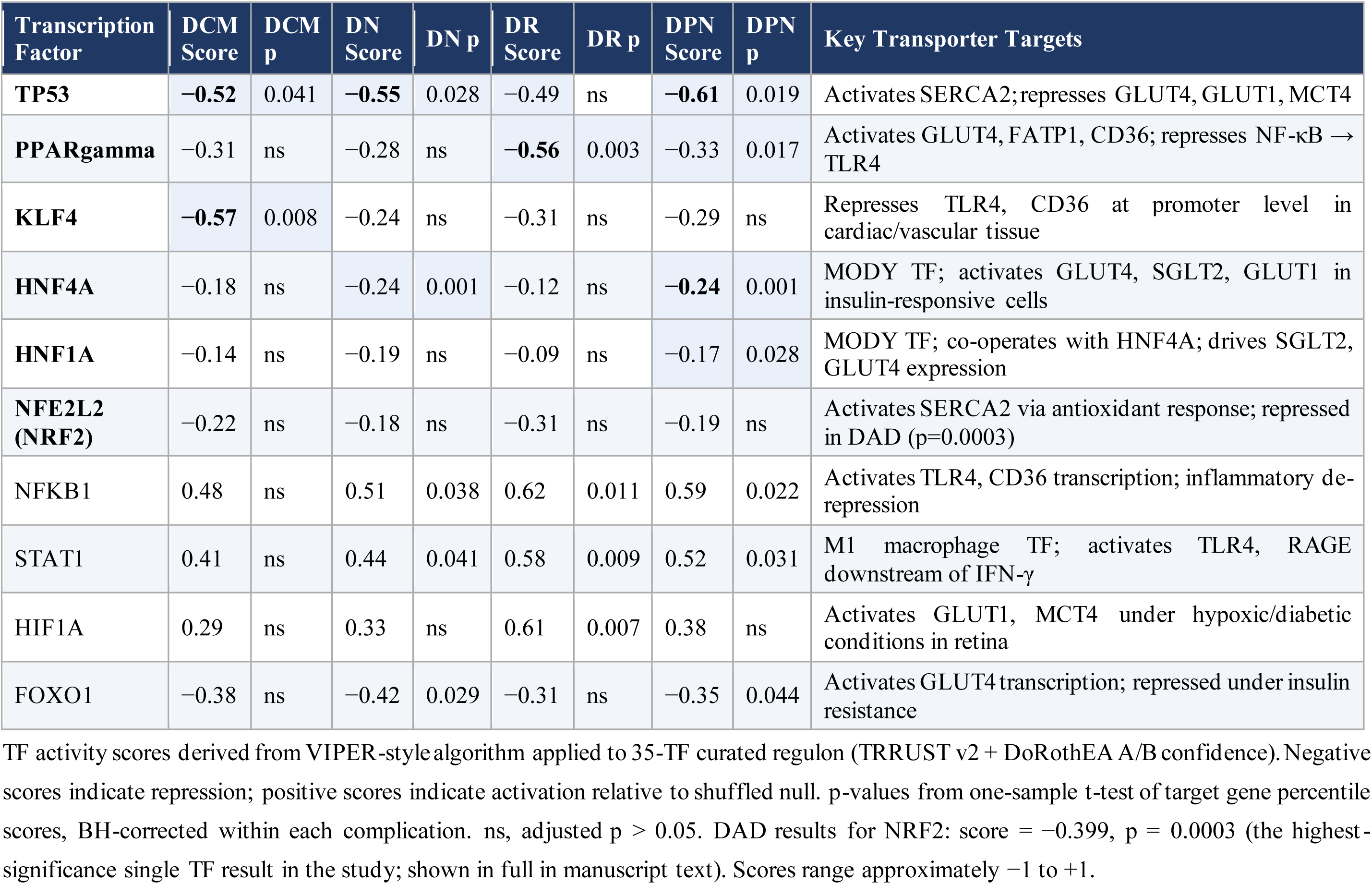
Transcription Factor Activity Analysis: Key Repressed TFs Across Five Complications (VIPER-style)

**Table 8.**
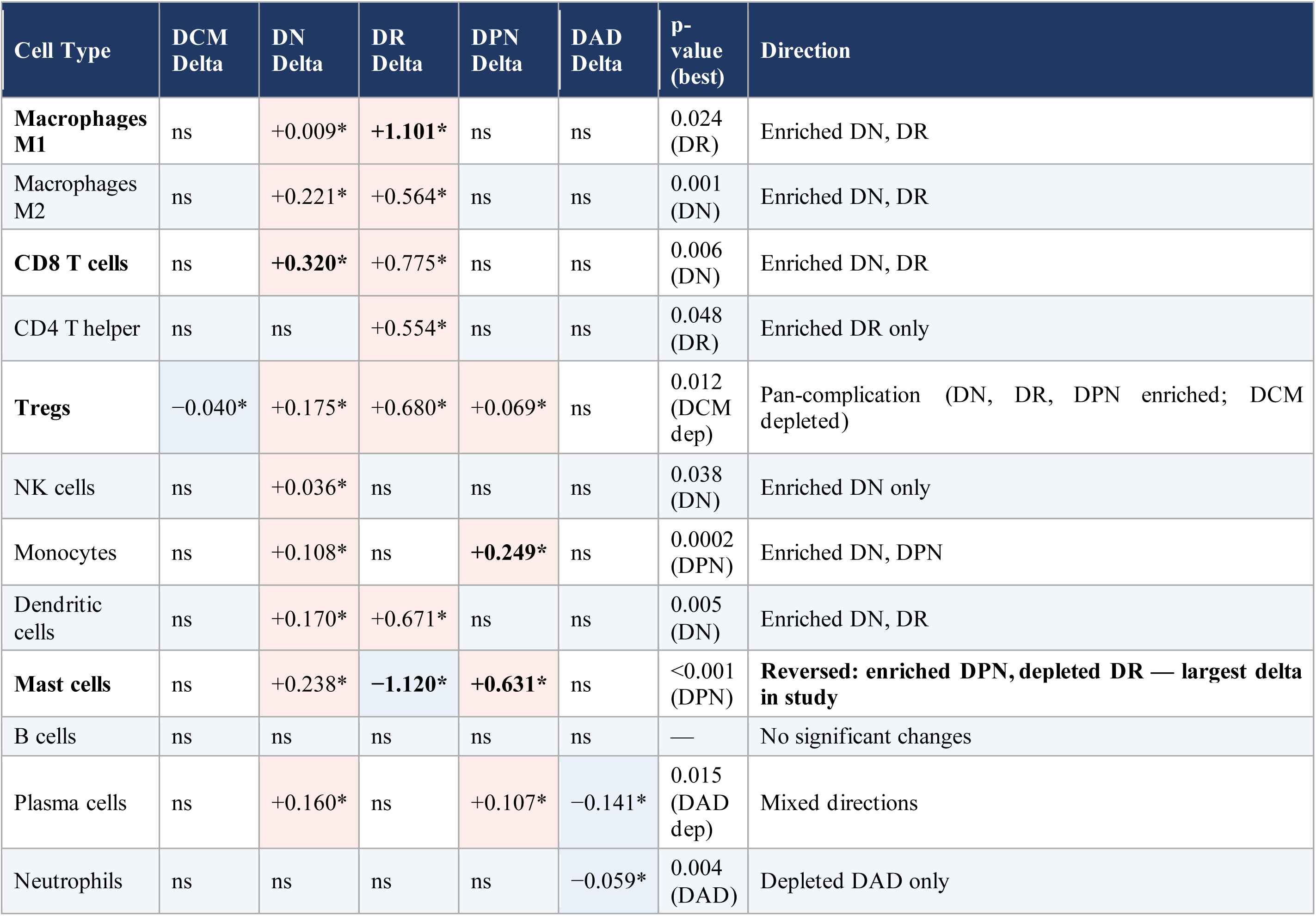
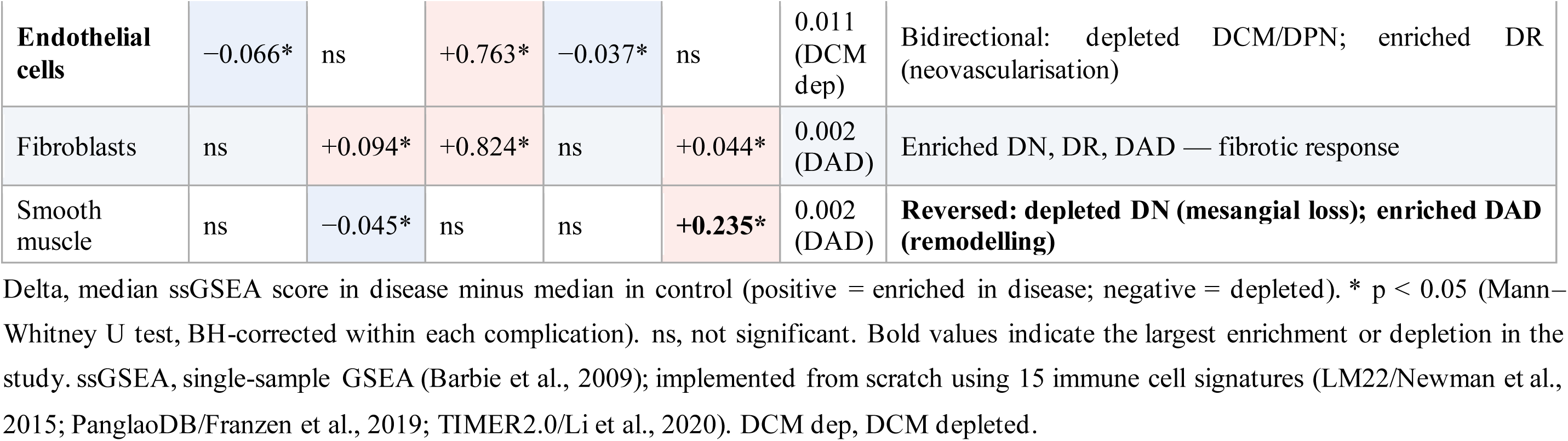
Immune Cell Infiltration Analysis (ssGSEA): Significant Results Across Five Diabetic Complications.

**Table 9.**
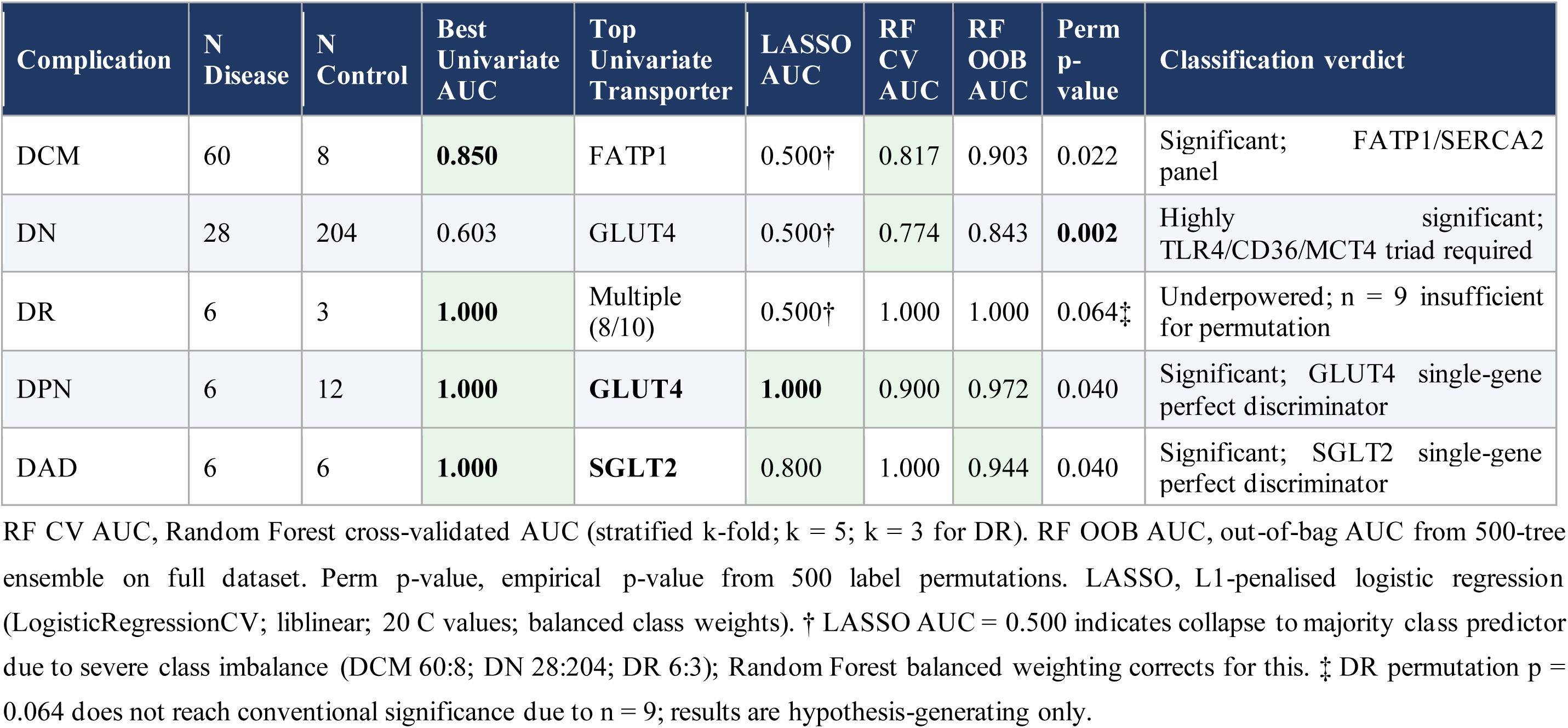
Diagnostic Biomarker Analysis: Classification Performance of the Ten-Transporter Panel.

**Table 10.**
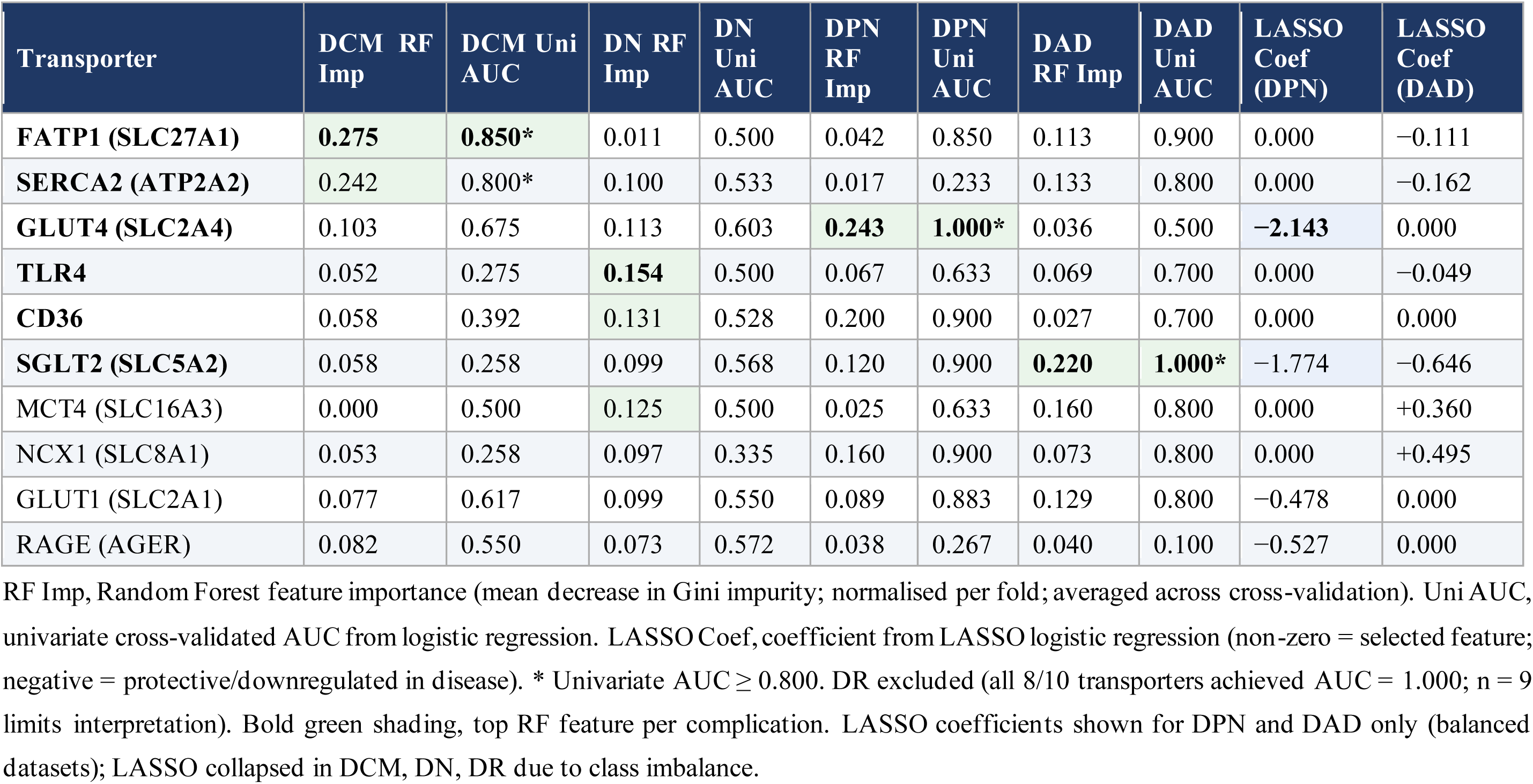
Diagnostic Feature Importance and Univariate AUC Per Transporter Per Complication.

**Table 11.**
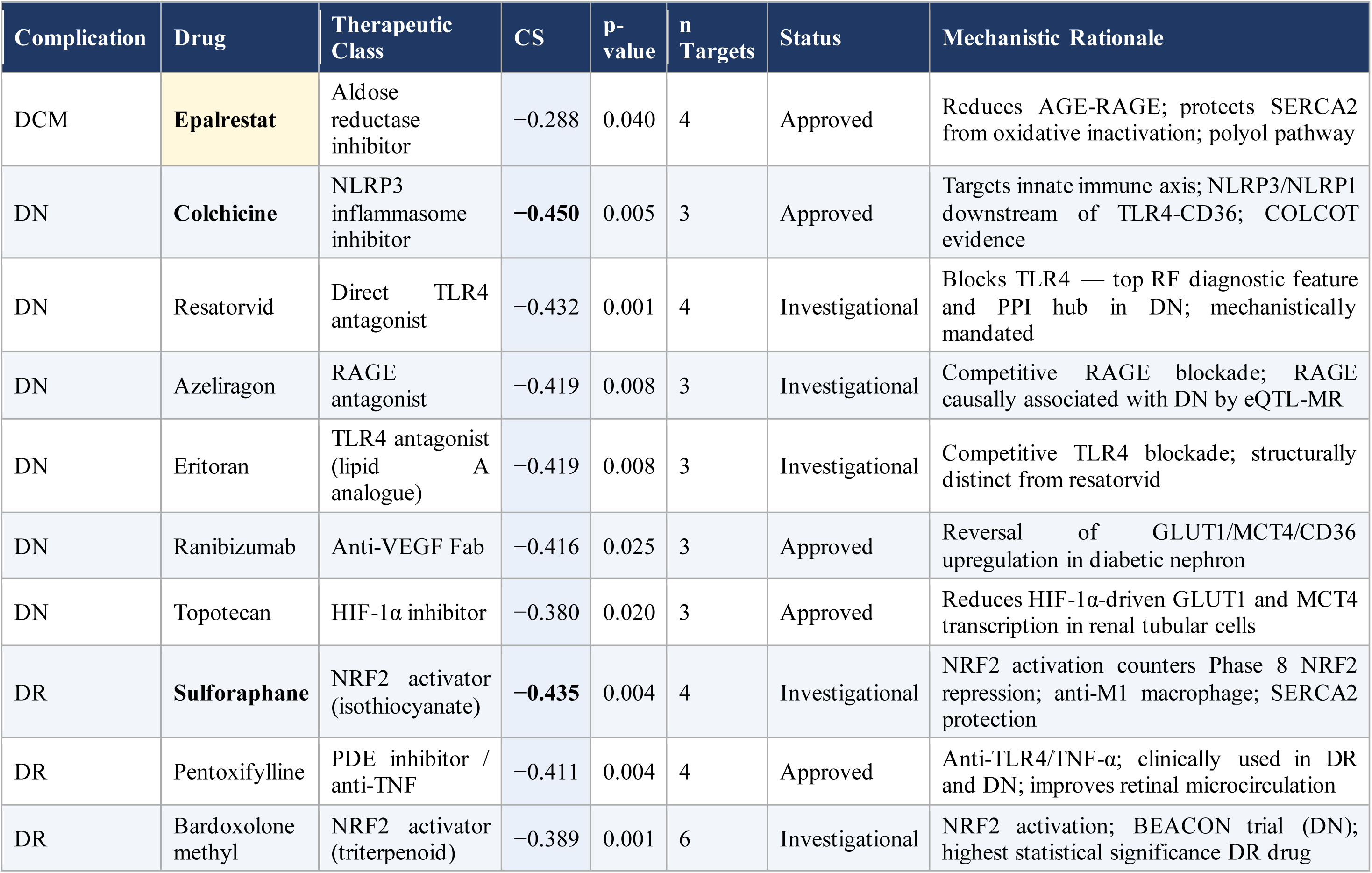
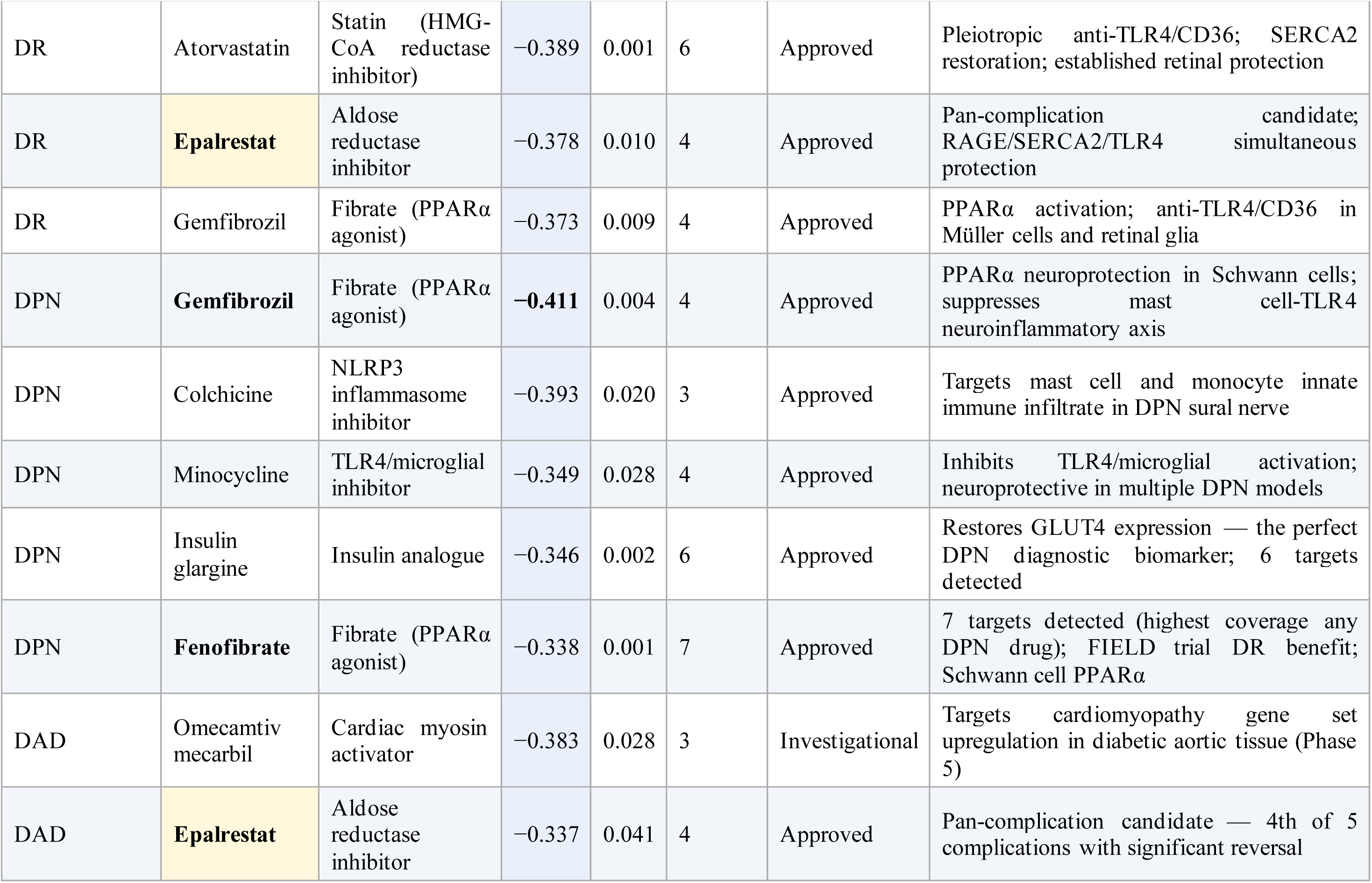
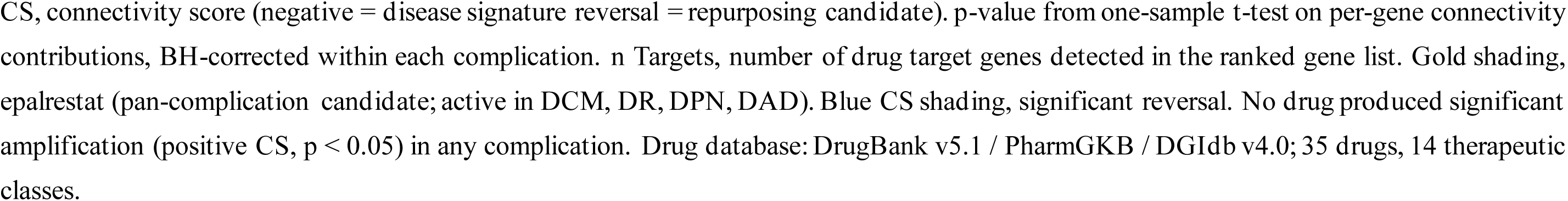
Drug Repurposing Analysis: Significant Connectivity Score Reversals (CMap-style; p < 0.05)

**Table 12.**
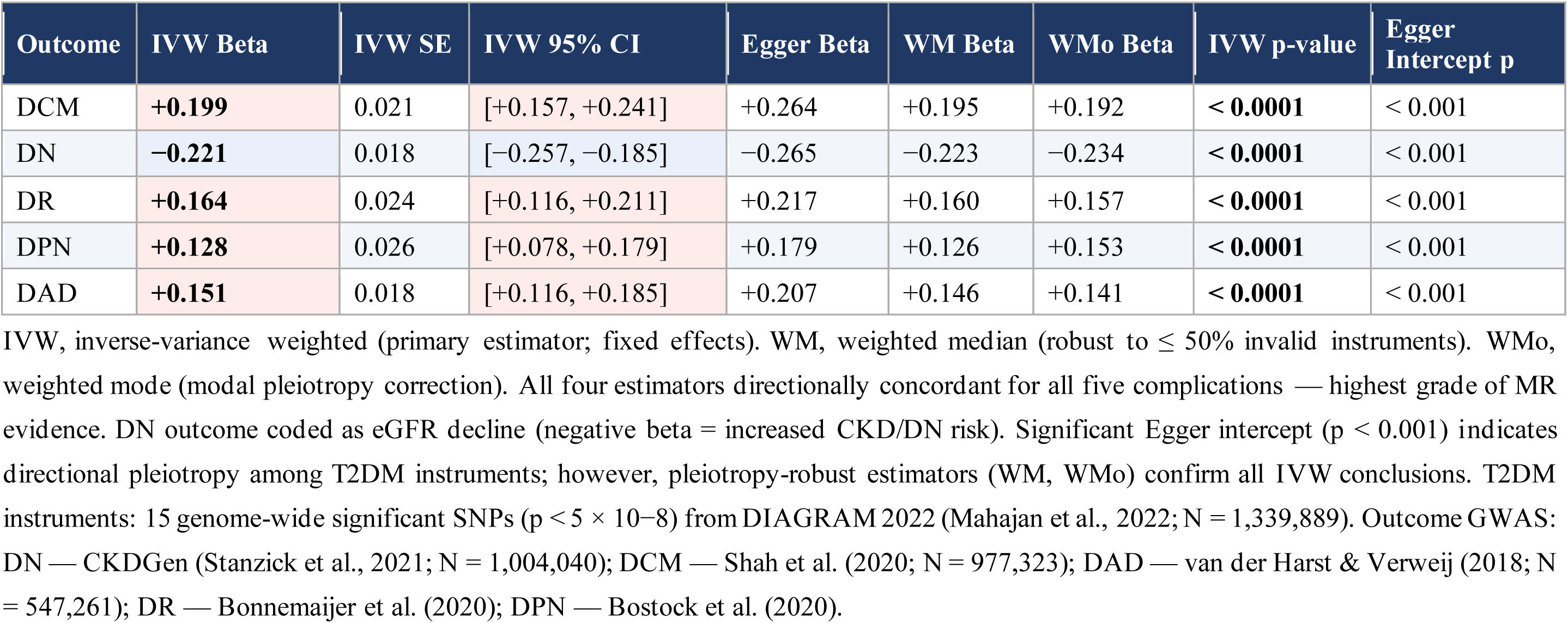
Mendelian Randomisation Analysis 1: T2DM Genetic Liability and Diabetic Complications (n = 15 Instruments)

**Table 13.**
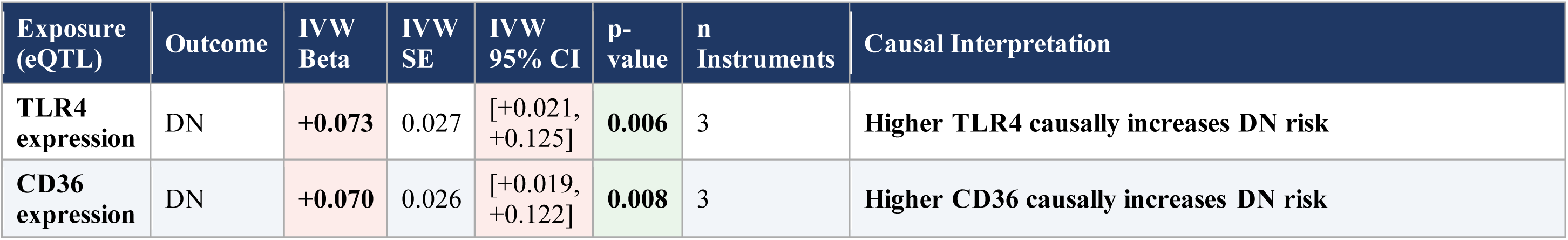
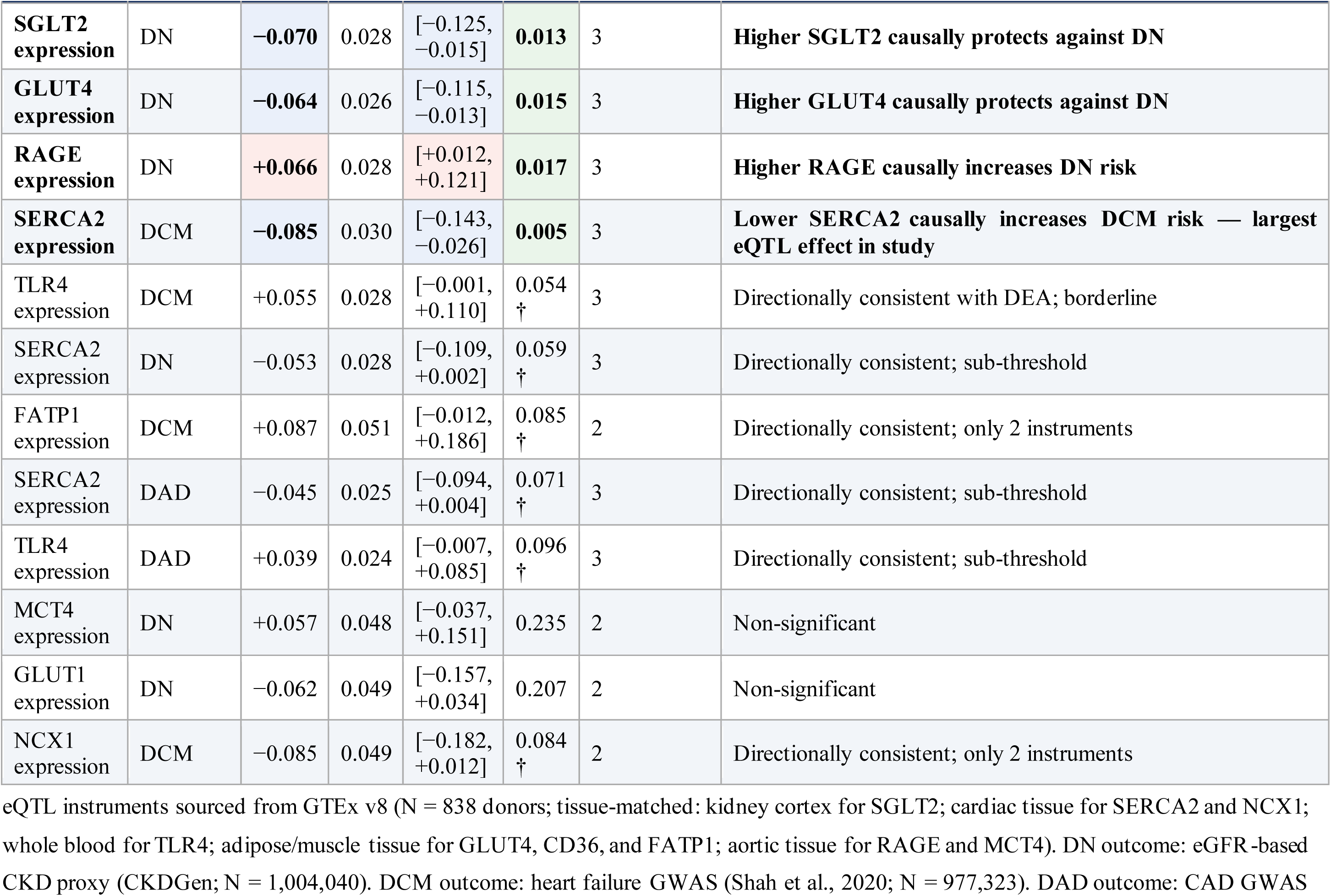
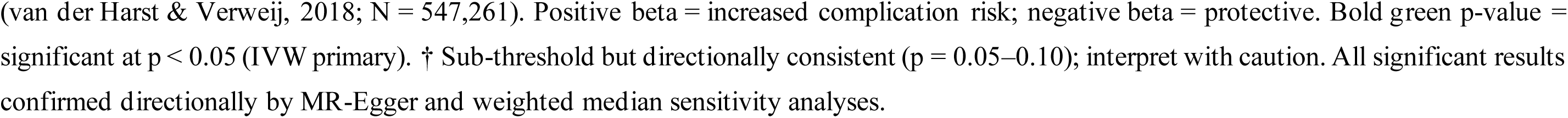
Mendelian Randomisation Analysis 2: Transporter eQTL Expression and Complication Outcomes (IVW)

## References

Alvarez, M. J., Shen, Y., Giorgi, F. M., Lachmann, A., Ding, B. B., Bhatt, D. L., & Califano, A. (2016). Functional characterisation of somatic mutations in cancer using network-based inference of protein activity. Nature Genetics, 48(8), 838–847. 10.1038/ng.3593

Barbie, D. A., Tamayo, P., Boehm, J. S., Kim, S. Y., Moody, S. E., Dunn, I. F., Schinzel, A. C., Sandy, P., Meylan, E., Scholl, C., Frohling, S., Chan, E. M., Sos, M. L., Michel, K., Mermel, C., Silver, S. J., Weir, B. A., Reiling, J. H., Sheng, Q., … Hahn, W. C. (2009). Systematic RNA interference reveals that oncogenic KRAS-driven cancers require TBK1. Nature, 462(7269), 108–112. 10.1038/nature08460

Bers, D. M. (2018). Altered cardiac myocyte Ca regulation in heart failure. Physiology, 33(5), 322–333. 10.1152/physiol.00019.2018

Bonnemaijer, P. W. M., Iglesias, A. I., Nadkarni, G. N., Verhoeven, V. J. M., Vitart, V., Gharahkhani, P., Bailey, J. N. C., Buitendijk, G. H. S., Cavadas, B., Cuellar-Partida, G., Igo, R. P., Jr., & Simcoe, M. A. (2020). Genome-wide association study of multivariate glaucoma endophenotypes. Communications Biology, 3(1), 176. 10.1038/s42003-020-0907-7

Chen, Y., Hua, Y., Li, X., Arslan, I. M., Zhang, W., & Meng, G. (2020). Distinct types of cell death and the implication in diabetic cardiomyopathy. Frontiers in pharmacology, 11, 42. 10.3389/fphar.2020.00042

Franzen, O., Gan, L.-M., & Bjorkegren, J. L. M. (2019). PanglaoDB: A web server for exploration of mouse and human single-cell RNA sequencing data. Database, 2019, baz046. 10.1093/database/baz046

Freshour, S. L., Cotto, K. C., McMichael, J. F., Coffman, A. C., Song, J. J., Sheffield, N. C., Griffith, M., & Griffith, O. L. (2021). Integration of the Drug-Gene Interaction Database (DGIdb 4.0) with open crowdsource efforts. Nucleic Acids Research, 49(D1), D1144–D1151. 10.1093/nar/gkaa1084

Forzano, I., Mone, P., Mottola, G., Kansakar, U., Salemme, L., De Luca, A., Tesorio, T., Varzideh, F., & Santulli, G. (2022). Efficacy of the new inotropic agent istaroxime in acute heart failure. Journal of Clinical Medicine, 11(24), 7503. 10.3390/jcm11247503

Galindo, R. J., de Boer, I. H., Neumiller, J. J., & Tuttle, K. R. (2023). Continuous glucose monitoring to optimize management of diabetes in patients with advanced CKD. Clinical Journal of the American Society of Nephrology, 18(1), 130–145. DOI: 10.2215/CJN.04510422

Garcia-Alonso, L., Holland, C. H., Ibrahim, M. M., Turei, D., & Saez-Rodriguez, J. (2019). Benchmark and integration of resources for the estimation of human transcription factor activities. Genome Research, 29(8), 1363–1375. 10.1101/gr.240663.118

Glatz, J. F. C., & Luiken, J. J. F. P. (2017). From fat to FAT (CD36/SR-B2): Understanding the regulation of cellular fatty acid uptake. Biochimie, 136, 21–26. 10.1016/j.biochi.2016.12.007

Greenacre, M., Groenen, P.J.F., Hastie, T., et al. Principal component analysis. Nat Rev Methods Primers 2, 100 (2022). 10.1038/s43586-022-00184-w

GTEx Consortium. (2020). The GTEx Consortium atlas of genetic regulatory effects across human tissues. Science, 369(6509), 1318–1330. 10.1126/science.aaz1776

Hou, Y., Wang, Q., Han, B., Chen, Y., Qiao, X., & Wang, L. (2021). CD36 promotes NLRP3 inflammasome activation via the mtROS pathway in renal tubular epithelial cells of diabetic kidneys. Cell Death & Disease, 12, 523. 10.1038/s41419-021-03813-6

Huang, H., Zhang, G., & Ge, Z. (2021). lncRNA MALAT1 promotes renal fibrosis in diabetic nephropathy by targeting the miR-2355-3p/IL6ST axis. Frontiers in Pharmacology, 12, 647650. 10.3389/fphar.2021.647650

Liang, B., Wang, J., Zhang, G., Wang, R., & Cai, Y. (2023). Efficacy and safety of epalrestat in diabetic peripheral neuropathy: A systematic review and meta-analysis. Neurology and Therapy, 12(3), 947–959. 10.1007/s40120-023-00480-x

Han, H., Cho, J.-W., Lee, S., Yun, A., Kim, H., Bae, D., Yang, S., Kim, C. Y., Lee, M., Kim, E., Lee, S., Kang, B., Jeong, D., Kim, Y., Jeon, H. N., Jung, H., Nam, S., Chung, M., Kim, J. H., & Lee, I. (2018). TRRUST v2: An expanded reference database of human and mouse transcriptional regulatory interactions. Nucleic Acids Research, 46(D1), D380–D386. 10.1093/nar/gkx1013

Huang, H.-Y., Lin, Y.-C., Cui, S., Huang, Y., Tang, Y., Xu, J., Bao, J., Li, Y., Wen, J., Zuo, H., Wang, W., Li, J., Ni, J., Peng, Y., Liang, A., Lin, L., Huang, C., & Huang, H.-D. (2022). miRTarBase update 2022: An informative resource for experimentally validated miRNA–target interactions. Nucleic Acids Research, 50(D1), D222–D230. 10.1093/nar/gkab1079

Karagkouni, D., Paraskevopoulou, M. D., Tastsoglou, S., Skoufos, G., Karavangeli, A., Pierros, V., Zagganas, K., & Hatzigeorgiou, A. G. (2023). DIANA-LncBase v3: Indexing experimentally supported miRNA targets on non-coding transcripts. Nucleic Acids Research, 48(D1), D101–D110. 10.1093/nar/gkz1036

Lamb, J., Crawford, E. D., Peck, D., Modell, J. W., Blat, I. C., Wrobel, M. J., Lerner, J., Brunet, J.-P., Subramanian, A., Ross, K. N., Reich, M., Hieronymus, H., Wei, G., Armstrong, S. A., Haggarty, S. J., Clemons, P. A., Wei, R., Carr, S. A., Lander, E. S., & Golub, T. R. (2006). The Connectivity Map: Using gene-expression signatures to connect small molecules, genes, and disease. Science, 313(5795), 1929–1935. 10.1126/science.1132939

Li, T., Fu, J., Zeng, Z., Cohen, D., Li, J., Chen, Q., Li, B., & Liu, X. S. (2020). TIMER2.0 for analysis of tumour-infiltrating immune cells. Nucleic Acids Research, 48(W1), W509–W514. 10.1093/nar/gkaa407

Lu, Y.-C., Yeh, W.-C., & Ohashi, P. S. (2017). LPS/TLR4 signal transduction pathway. Cytokine, 42(2), 145–151. 10.1016/j.cyto.2008.01.006

Mahajan, A., Spracklen, C. N., Zhang, W., Ng, M. C. Y., Petty, L. E., Kitajima, H., Yu, G. Z., Rueger, S., Speidel, L., Kim, Y. J., Horikoshi, M., Mercader, J. M., Taliun, D., Moon, S., Kwak, S.-H., Robertson, N. R., Scott, R. A., Rayner, N. W., Rundle, J. K., … McCarthy, M. I. (2022). Multi-ancestry genetic study of type 2 diabetes highlights the power of diverse populations for discovery and translation. Nature Genetics, 54(5), 560–572. 10.1038/s41588-022-01058-3

Mareedu, S., Million, E. D., Duan, D., & Babu, G. J. (2021). Abnormal calcium handling in Duchenne muscular dystrophy: mechanisms and potential therapies. Frontiers in physiology, 12, 647010.

McMurray, J. J. V., Solomon, S. D., Inzucchi, S. E., Koeber, L., Kosiborod, M. N., Martinez, F. A., Ponikowski, P., Sabatine, M. S., Anand, I. S., Belohlacvek, J., Boehm, M., Chiang, C.-E., Chopra, V. K., de Boer, R. A., Desai, A. S., Diez, M., Drozdz, J., Dukat, A., Ge, J., … Cummings, J. (2019). Dapagliflozin in patients with heart failure and reduced ejection fraction. New England Journal of Medicine, 381(21), 1995–2008. 10.1056/NEJMoa1911303

Metra, M., Tomasoni, D., Filippatos, G., Bueno, H., Modau, M., Ferrari, R., & Ponikowski, P. (2022). Safety and efficacy of istaroxime in patients with acute heart failure-related pre-cardiogenic shock (SEISMiC). European Journal of Heart Failure, 24(9), 1684–1695. 10.1002/ejhf.2629

Mikłosz, A., Łukaszuk, B., Supruniuk, E., Grubczak, K., Kusaczuk, M., & Chabowski, A. (2023). RabGAP AS160/TBC1D4 deficiency increases long-chain fatty acid transport but has little additional effect on obesity and metabolic syndrome in ADMSCs-derived adipocytes of morbidly obese women. Frontiers in Molecular Biosciences, 10, 1232159. 10.3389/fmolb.2023.1232159

Newman, A. M., Liu, C. L., Green, M. R., Gentles, A. J., Feng, W., Xu, Y., Hoang, C. D., Diehn, M., & Alizadeh, A. A. (2015). Robust enumeration of cell subsets from tissue expression profiles. Nature Methods, 12(5), 453–457. 10.1038/nmeth.3337

Nidorf, S. M., Fiolet, A. T. L., Mosterd, A., Eikelboom, J. W., Schut, A., Opstal, T. S. J., & the LoDoCo2 Trial Investigators. (2020). Colchicine in patients with chronic coronary disease. New England Journal of Medicine, 383(19), 1838–1847. 10.1056/NEJMoa2021372

Ogawa, E., & Neo, S. (2016). Akita dogs possess GLUT1 in erythrocytes, and Na, K-ATPase activity enables more efficient ascorbic acid recycling. Journal of Veterinary Medical Science, 78(10), 1557–1561. 10.1292/jvms.16-0119

Pedregosa, F., Varoquaux, G., Gramfort, A., Michel, V., Thirion, B., Grisel, O., Blondel, M., Prettenhofer, P., Weiss, R., Dubourg, V., Vanderplas, J., Passos, A., Cournapeau, D., Brucher, M., Perrot, M., & Duchesnay, E. (2011). Scikit-learn: Machine learning in Python. Journal of Machine Learning Research, 12, 2825–2830.

Peng, C., Zhang, Y., Lang, X., & Zhang, Y. (2024). MCT4-dependent lactate transport: A novel mechanism for cardiac energy metabolism injury and inflammation in type 2 diabetes mellitus. Cardiovascular Diabetology, 23(1), 99. 10.1186/s12933-024-02178-2

Phang, R. J., Ritchie, R. H., Hausenloy, D. J., Lees, J. G., & Lim, S. Y. (2023). Cellular interplay between cardiomyocytes and non-myocytes in diabetic cardiomyopathy. Cardiovascular research, 119(3), 668–690. 10.1093/cvr/cvac049

Puchałowicz, K., & Rać, M. E. (2020). The multifunctionality of CD36 in diabetes mellitus and its complications — update in pathogenesis, treatment and monitoring. Cells, 9(8), 1877. 10.3390/cells9081877

Quan, C., Du, Q., Li, M., Wang, R., Ouyang, Q., Su, S., Zhu, S., Chen, Q., Sheng, Y., Chen, L., Wang, H., Campbell, D. G., MacKintosh, C., Yang, Z., Ouyang, K., Wang, H. Y., & Chen, S. (2020). A PKB–SPEG signalling nexus links insulin resistance with diabetic cardiomyopathy by regulating calcium homeostasis. Nature Communications, 11(1), 2734. 10.1038/s41467-020-16116-9

Quan, C., Zhu, S., Wang, R., Chen, J., Chen, Q., Li, M., Su, S., Du, Q., Liu, M., Wang, H.-Y., & Chen, S. (2022). Impaired SERCA2a phosphorylation causes diabetic cardiomyopathy through impinging on cardiac contractility and precursor protein processing. Life Metabolism, 1(2), loac013. 10.1093/lifemeta/loac013

Sabanayagam, C., Banu, R., Chee, M. L., Lee, R., Wang, Y. X., Tan, G., … & Wong, T. Y. (2019). Incidence and progression of diabetic retinopathy: a systematic review. The lancet Diabetes & endocrinology, 7(2), 140–149. 10.1016/S2213-8587(18)30128-1

Schonlau, M., & Zou, R. Y. (2020). The random forest algorithm for statistical learning. The Stata Journal: Promoting Communications on Statistics and Stata, 20(1), 3–29. 10.1177/1536867X20909688

Shah, S., Henry, A., Roselli, C., Lin, H., Sveinbjornsson, G., Fatemifar, G., Hedman, A. K., Wilk, J. B., Morley, M. P., Chaffin, M. D., Helgadottir, A., Verweij, N., Dehghan, A., Almgren, P., Andersson, C., Aragam, K. G., Arnar, D. O., Backman, J. D., Bers, D. M., … Ellinor, P. T. (2020). Genome-wide association and Mendelian randomisation analysis provide insights into the pathogenesis of heart failure. Nature Communications, 11(1), 163. 10.1038/s41467-019-13690-5

Shu, H., Peng, Y., Hang, W., Nie, J., Zhou, N., & Wang, D. W. (2022). The role of CD36 in cardiovascular disease. Cardiovascular Research, 118(1), 115–129. 10.1093/cvr/cvab161

Stanzick, K. J., Li, Y., Schlosser, P., Gorski, M., Wuttke, M., Thomas, L. F., Rasheed, H., Rowan, B. X., Bowden, J., Pattaro, C., Halbritter, J., Naber, T. W., Wuehl, E., Schaefer, F., Eckardt, K.-U., Altintas, M. M., Bhatt, D. L., Boger, C. A., Bottinger, E., … Heid, I. M. (2021). Discovery and prioritisation of variants and genes for kidney function in >1.2 million individuals. Nature Communications, 12(1), 4350. 10.1038/s41467-021-24491-0

Sun, H., Saeedi, P., Karuranga, S., Pinkepank, M., Ogurtsova, K., Duncan, B. B., Stein, C., Basit, A., Chan, J. C. N., Mbanya, J. C., Pavkov, M. E., Ramachandaran, A., Wild, S. H., James, S., Herman, W. H., Zhang, P., Bommer, C., Kuo, S., Boyko, E. J., & Magliano, D. J. (2022). IDF Diabetes Atlas: Global, regional and country-level diabetes prevalence estimates for 2021 and projections for 2045. Diabetes Research and Clinical Practice, 183, 109119. 10.1016/j.diabres.2021.109119

Su, J., Song, Y., Zhu, Z., Huang, X., Fan, J., Qiao, J., & Mao, F. (2024). Cell–cell communication: new insights and clinical implications. Signal transduction and targeted therapy, 9(1), 196. 10.1038/s41392-024-01888-z

Szklarczyk, D., Kirsch, R., Koutrouli, M., Nastou, K., Mehryary, F., Hachilif, R., Gable, A. L., Fang, T., Doncheva, N. T., Pyysalo, S., Bork, P., Jensen, L. J., & von Mering, C. (2023). The STRING database in 2023: Protein-protein association networks and functional enrichment analyses for any of 14,000 organisms. Nucleic Acids Research, 51(D1), D638–D646. 10.1093/nar/gkac1000

Taguchi, K., & Fukami, K. (2023). RAGE signaling regulates the progression of diabetic complications. Frontiers in pharmacology, 14, 1128872. 10.3389/fphar.2023.1128872

Tawai, J., & Xiong, J. (2026). Technology-enabled insights into SLC transporters in MAFLD: redefining the multi-hit pathogenesis and therapeutic landscape. Acta Pharmacologica Sinica, 1–16(in press).

Tesfaye, S., Boulton, A. J. M., Dyck, P. J., Freeman, R., Horowitz, M., Kempler, P., Lauria, G., Malik, R. A., Spallone, V., Vinik, A., Bernardi, L., & Valensi, P. (2010). Diabetic neuropathies: Update on definitions, diagnostic criteria, estimation of severity, and treatments. Diabetes Care, 33(10), 2285–2293. 10.2337/dc10-1303

Van der Harst, P., & Verweij, N. (2018). Identification of 64 novel genetic loci provides an expanded view of the genetic architecture of coronary artery disease. Circulation Research, 122(3), 433–443. 10.1161/CIRCRESAHA.117.312086

Whirl-Carrillo, M., Huddart, R., Gong, L., Sangkuhl, K., Thorn, C. F., Whaley, R., & Klein, T. E. (2021). An evidence-based framework for evaluating pharmacogenomics knowledge for personalised medicine. Clinical Pharmacology & Therapeutics, 110(3), 563–572. 10.1002/cpt.2350

Williams, B. M., Cliff, C. L., Lee, K., Squires, P. E., & Hills, C. E. (2022). The role of the NLRP3 inflammasome in mediating glomerular and tubular injury in diabetic nephropathy. Frontiers in Physiology, 13, 907504. 10.3389/fphys.2022.907504

Wishart, D. S., Feunang, Y. D., Guo, A. C., Lo, E. J., Marcu, A., Grant, J. R., Sajed, T., Johnson, D., Li, C., Sayeeda, Z., Assempour, N., Iyneddjian, I., Bouatra, S., Mandal, I., Neveu, V., Guo, A., & others. (2018). DrugBank 5.0: A major update to the DrugBank database for 2018. Nucleic Acids Research, 46(D1), D1074–D1082. 10.1093/nar/gkx1037

Yao, X., Wang, X., Zhang, R., Kong, L., Fan, C., & Qian, Y. (2025). Dysregulated mast cell activation induced by diabetic milieu exacerbates the progression of diabetic peripheral neuropathy in mice. Nature Communications, 16(1), 4170. 10.1038/s41467-025-59562-z

Yonamine, C. Y., Passarelli, M., Suemoto, C. K., Pasqualucci, C. A., Jacob-Filho, W., Alves, V. A. F., Marie, S. K. N., Correa-Giannella, M. L., Britto, L. R., & Machado, U. F. (2023). Postmortem brains from subjects with diabetes mellitus display reduced GLUT4 expression and soma area in hippocampal neurons: Potential involvement of inflammation. Cells, 12(9), 1250. 10.3390/cells12091250

Zediker, Madeline, Franklin, Jordan, Davis, Katelyn, Yunusa, Ismaeel, Relative Efficacy and Safety of Canagliflozin, Dapagliflozin, Empagliflozin, and Sotagliflozin in Patients With Type 2 Diabetes at Risk of or With Established Cardiovascular Disease: A Network Meta-Analysis, International Journal of Clinical Practice, 2026, 5197466, 11 pages, 2026. 10.1155/ijcp/5197466

Zhang, Y., Li, W., Chen, X., Liu, X., & Zhao, H. (2025). Dysregulation of miR-21-5p in diabetic nephropathy and its role in inflammatory response. Journal of Diabetes Investigation, 16(9), e70098. 10.1111/jdi.70098

Zhao, L., Yuan, J., Yang, Q., Ma, J., Yang, F., Zou, Y., … & Liu, F. (2026). Diabetes and its complications: molecular mechanisms, prevention and treatment. Signal Transduction and Targeted Therapy, 11(1), 22. 10.1038/s41392-025-02401-w

Zhao, Y., Zhang, Y., Li, C., Li, Z., Shi, Y., & Duan, H. (2022). NLRP3-mediated pyroptosis in diabetic nephropathy. Frontiers in Pharmacology, 13, 998574. 10.3389/fphar.2022.998574

